# Stem-Cell-Derived Islets as a Model of Human Islet Inflammation: A Comparative Analysis of Pro-inflammatory Cytokine Responses

**DOI:** 10.64898/2026.05.01.722128

**Authors:** Cecilie Amalie Brøgger Svane, Adam Bøgh Marstrand-Jørgensen, Anne Jørgensen, Rebekka Gerwig, Josefine Gudman, Tina Fløyel, Tarun Veer Singh Ahluwalia, Flemming Pociot, Joachim Størling

## Abstract

**Background:** Inflammation-induced pancreatic islet-cell death and dysfunction are key aspects of both type 1 and type 2 diabetes. Stem cell-derived islets (SC-islets) are an emerging tool in diabetes research, however, our understanding of how inflammation affects SC-islet function is incomplete. We therefore aimed to thoroughly characterize how SC-islets respond to pro-inflammatory cytokines at the functional and transcriptomic levels in comparison with human primary islets and EndoC-βH5 cells.

**Method:** A 7-stage differentiation protocol was used to generate SC-islets with insulin-, glucagon-, and somatostatin-positive cells. SC-islets, primary human islets and EndoC-βH5 cells were exposed to different doses of pro-inflammatory cytokines (IL-1β + IFNγ + TNFα) including a high dose for up to 48 h and a low dose up to 144 h to mimic the intense islet inflammation in T1D and chronic low-grade inflammation in T2D, respectively. Differential gene expression (RNA-seq), cell death, activation of key signalling proteins, hormones, and chemokine secretion were determined.

**Results:** Basal expression of key islet-cell identity genes in SC-islets correlated well with that of primary islets and EndoC-βH5 cells. In SC islets, cytokines dose-dependently induced activation of key proximal signalling pathways (NFκB, STAT1, and JNK), upregulation of major histocompatibility complex (MHC) class I, and increased cell death (cytotoxicity and caspase 3/7 activity). In head-to-head experiments, SC-islets displayed similar cytokine responses particularly as primary islets regarding induction of cell death, chemokine secretion, differential gene expression, and protein levels of cell death executioners (gasdermin D and caspase-7). Cytokines increased insulin release in SC-islets and primary islets, while diminishing insulin secretion in EndoC-βH5 cells. Cytokines reduced glucagon release in SC-islets, which was partially restored by treatment with the incretin hormone glucose-dependent insulinotropic peptide (GIP) with or without a glucagon-like peptide 1 (GLP-1) receptor agonist (liraglutide).

**Conclusion:** SC-islets are highly responsive to inflammation with a high degree of similarity to primary islets. Our results support the use of SC-islets as a valid tool in inflammation and diabetes research.

## Introduction

The endocrine pancreatic islet cells – the beta-, alpha- and delta cells – are responsible for the tight regulation of blood glucose levels [1]. In both type 1 (T1D) and type 2 diabetes (T2D), local inflammation within the islets plays a prominent role in disease progression by causing functional impairment and death of the beta cells leading to absolute or partial insulin deficiency [2]. Consequently, the inflammation-induced islet-cell demise has profound effects on glucose homeostasis [3, 4]. Pro-inflammatory cytokines are key contributors to islet inflammation and exposure of islets to cytokines in vitro is a widely used model for mimicking the inflammatory insult during diabetes pathogenesis. The classical diabetogenic cytokines used are interleukin (IL)-1β, interferon (IFN)-γ, and tumour necrosis factor (TNF)-α [5]. These cytokines are secreted by activated islets-resident dendritic cells and macrophages and by the infiltrating T lymphocytes, which have been shown to recapitulate the pro-apoptotic and toxic microenvironment of islet inflammation [6]. Interestingly, the endocrine islets cells are also a source of inflammatory factors as increased cytokine and chemokine expression and secretion are seen in cytokine-exposed islets and beta cells [7–9]. Human islets proportionally contain more non-beta cells compared to rodent islets [10], which may explain species differences in inflammatory responses and sensitivity to cytokines [11]. The availability of isolated primary human islets for research is limited and in combination with a high degree of heterogeneity between donors and batches, human islets is challenging model system [1, 12]. A few human beta-cell lines have been developed from pancreas tissue of human foetuses including the EndoC-βH5 cells [13]. We recently characterized these cells’ sensitivity and responsiveness to inflammation (proinflammatory cytokines) validating them as a tool for studying inflammation-mediated beta-cell dysfunction [8]. However, the use of a single endocrine cell type, i.e., beta cells, fails to satisfactory model the complex interplay between different islet-cell types and is therefore not an optimal model system for human islet research.

In vitro differentiation of pluripotent stem cells has emerged as a promising tool to generate islet-like aggregates (SC-islets) for disease modelling studies [14–19]. While SC-islets share many similarities with primary islets, further characterization of SC-islets and their response to inflammatory conditions is warranted to validate their translational use in islet inflammation research. To date, only a few studies have investigated the responsiveness of SC-islets to pro-inflammatory cytokines, i.e. IL-1β, IFNγ, TNF-α, and IFNα [20–22]. In these studies, cytokine treatment of SC-islets induced expression of relevant genes and proteins, e.g., HLA-A/B/C and markers for endoplasmic reticulum (ER) stress, led to supressed hormone expression, increased chemokine C-X-C motif chemokine Ligand (CXCL)-10 secretion, and induced apoptosis. Additionally, SC-islets are a promising tool for studying candidate risk genes influence on development and pathogenesis, in which the characterization of their basal response is essential. Hence, SC-islets hold promise as a model system harbouring the cell composition and complexity as native islets, however, their use in islet inflammation research is still in the early phase warranting additional studies to uncover their full potential in this field.

In this study, we generated SC-islets and thoroughly characterized their responses to proinflammatory cytokines including direct comparative analyses with isolated primary human islets and EndoC-βH5 cells. We assessed the impact of different concentrations of cytokines (IL-1β, IFNγ and TNFα) for up to 144 h to mimic the clinical conditions of T1D and T2D, i.e., ‘acute’ high- and chronic low-grade inflammation [8, 23]. Overall, our findings demonstrate good consistency between the model systems particularly between SC-islets and primary islets, supporting SC-islets as a valid model for islet inflammation research.

## Methods/materials

### Cell culture

Human beta-cell line EndoC-βH5 cells (Human Cell Design, France) were maintained as previously described [8]. EndoC-βH5 were seeded at 100.000 cells/well in 96-well plates coated with β-coat (Human Cell Design). EndoC-βH5 were maintained in Ultiβ1 culture medium (Human Cell Design). Human pancreatic primary islets from non-diabetic donors were obtained through Prodo Laboratories Inc. (USA). Donor details are listed in Supplementary Table 1. The primary islets were cultured in Ham’s F10 nutrient mix (Gibco), supplemented with 10% heat inactivated foetal bovine serum (hiFBS) and 100 U/mL Penicillin and 100 ug/mL Streptomycin during, which during experiments was replaced with 2% human serum (Sigma) and 10 U/mL Penicillin and 10 ug/mL Streptomycin. The Human Induced Pluripotent Stem cell (IPSC) line 1023 was kindly provided by Associate Professor Dietrich M. Egli, University of Columbia. Expression of pluripotency markers, germ layer differentiation and Karyotype (performed at the Kennedy Center Glostrup, DK) were validated prior to initiation of the study (Supplementary Figure 1). Mycoplasma level was confirmed negative using Mycoplasma Alert detection kit (Lonza, LT07). The IPSCs were cultured on Matrigel (Corning)-coated plates in mTeSR^TM^ Plus medium (Stemcell Technologies) and passaged every 5-8 days using 0.5 μM EDTA in PBS. On day -2, mTeSR^TM^ Plus medium was replaced with Essential 8 (E8) medium (Gibco). The differentiation was carried out using a seven-stage differentiation protocol adapted from [16, 18, 24]. Media components and supplements are listed in Supplementary Table 2. Briefly, IPSCs were passaged in Accutase (Stemcell Technologies) on day -1 and seeded at 2.25e5 cells/cm^2^ in E8 with 10 μM Rho-associated Kinase inhibitor Y-27632 (ROCKi) (Millipore and Stem Cell Technologies). From day 0, complete daily media change was performed according to Supplementary Table 2. On day 12, the endocrine progenitor cells were passaged in Accutase and seeded in non-adherent microwells (Stemcell Technology) with 750 cells/microwell in stage 5 media with 10 μM ROCKi and 20 ng/mL Heparin (Sigma). Daily 50% media changes were performed during stage 5. At stage 6, the aggregates were transferred to suspension and placed on orbital shaker, with a 50% media change every 2-3 days until day 45. Stage validations were performed at end stage 1, 4, and end-differentiation (representative images in Figure 1B and Supplementary Figure 2).

**Figure 1.**
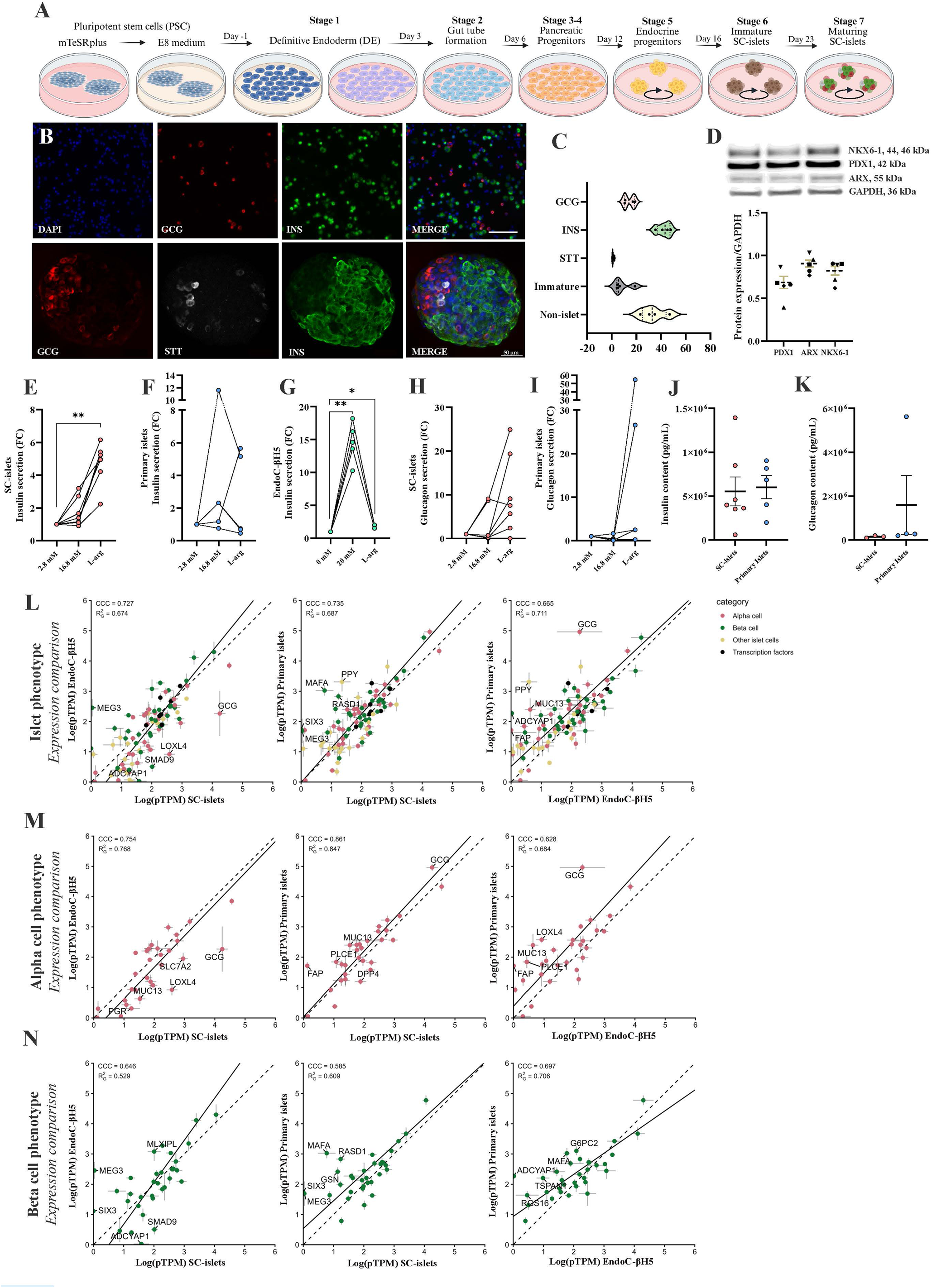
SC-islets characterization with comparison to primary human islets and human EndoC-βH5 cells. **A)** Differentiation process following 7-stage protocol, created in biorender. **B)** Representative immunofluorescent images of single-celled SC-islets (top, scalebar: 200 nm) and whole SC-islets (bottom, scalebar: 50 nm), with Insulin (green), Glucagon (red) and somatostatin (white) and nuclei (blue). **C)** IF staining quantification of SC-islet hormone expression (%). **D)** Western blot protein levels of PDX1, NKX6-1 and ARX normalized to GAPDH. Fold change (FC) hormone secretion at low glucose (2.8 mM), high glucose (16.8 mM) and 15 mM L-arginine in low glucose, measured for insulin in **E)** SC-islets (N = 7) **F)** Primary islets (N = 5) and **G)** EndoC-βH5 cells (N = 3-5) and glucagon in **H)** SC-islets (N = 7) and **I)** primary islets (N = 5). Islet hormone content for **J)** insulin and **K)** glucagon from SC-islets (N = 3-7) and Primary islets (N = 4-5). Graphs (C-K) presented as mean±SEM, with individual data points for biological replicate or donor. Paired t-test performed for graphs E-K, with data considered significant with p-value <0.05; *<0.05, **<0.01. Log10(pTPM+1) Deming regression correlation of all models for **L)** islet-identity genes and identity stratified for **M)** alpha- and **N)** beta-cell specific genes. H-J striped lines depict an exact correspondence, with solid line as actual Deming regression. CCC = correspondence correlation coefficient for level of expression and R2g = Generalized R Squared for linear regression of expression pattern.

### Cytokine stimulation

Cytokine dose response was performed by titration of 1000 U/mL recombinant human (rh) TNFα (R&D systems), 1000 U/mL rhIFNγ (Peprotech) and 50 U/mL rhIL-1β (R&D systems). The doses were selected based on published investigations for induction of beta- and alpha-cell dysfunction [8, 23].

### Hormone secretion assay

For the EndoC-βH5, a complete media change to Ulti-Starvation (-ST) (Human Cell Design) medium was performed 24h prior to assay. The assay was performed as previously described [8], with 0 mM glucose, 20 mM glucose and 15 mM L-Arginine (Sigma) in 0 mM, followed by cell lysis for content. Sequential GSIS was performed on islet models. Following 90-minute reset in 2.8 mM glucose (sigma) in Krebs Ringer Buffer (0.2% fatty acid free foetal bovine serum (fafBSA) (sigma), 0.04% NaHCO_3_ (Gibco), and 1 mM HEPES (Gibco)), islets were incubated for 30-minute intervals in 2.8 mM glucose, 16.8 mM glucose and 15 mM L-Arginine in 2.8 mM glucose, followed by cell lysis for content. Incretin-regulated secretion was performed by 1 μM Glucagon-Like Peptide-1 (GLP-1) agonist Liraglutide (BioTechne), 1 μM Glucose-Dependent Insulinotropic peptide (GIP) (BioTechne) or in combination, supplemented from reset. Insulin and glucagon were measured with Human Insulin ELISA and Human Glucagon ELISA (Mercodia). Absorbance was read on Infinite M200 PRO plate reader (Tecan, Switzerland), software Magellan 7.2. Glucagon was additionally measured with U-PLEX Human Glucagon Assay (K1515YK, Mesoscale discovery (MSD)), electrochemiluminescence read on MESO QuickPlex SQ 120MM.

### Immunofluorescence and image analysis

Whole aggregates, monolayer or dispersed SC-islets were fixed for 15-60 minutes (monolayer/aggregates) in 4% PFA in PBS (ThermoFisher), permeabilized in 0.5% TritonX100 (Sigma) followed by blocking in 5% Donkey Serum (Jacksons ImmunoResearch). Cells were incubated with primary antibodies, diluted in 5% Donkey Serum with 0.1% tween20 (Sigma) for 2h RT or 4°C O/N, followed by secondary antibodies for 1h, RT. Antibody and dilutions are listed in Supplementary Table 3. Nucleus staining was performed with Vectashield with DAPI (Vector Laboratories) or 1:1000 DAPI (Sigma) in PBS.

ImageXpress PICO Automated Cell Imaging System (Molecular Devices) was used to image with quantification in CellReporterXpress 2.9.3, with protocol *cell scoring* for two markers. Representative images of checkpoints were visualized using the Nikon ECLIPSE Ti2 microscope (Nikon). For confocal microscopy, whole aggregates were stained following the described protocol and mounted on microscope slides with Prolong Gold antifade mounting medium (ThermoFisher). Zeiss Imager.M2 LMS900 confocal microscope with ZEN 3.3 system software was used to visualize the cells.

### Cell death assays

The Caspase-Glo® 3/7 Assay (Promega) and CytoTox-Glo^TM^ Cytotoxicity Assay (Promega) were performed according to manufacturer protocols and measured for luminescence on Infinite M200 PRO plate reader (Tecan) with Magellan 7.2 software.

### Chemokine secretion

Secreted chemokines were measured on V-PLEX Chemokine panel 1 Human (K15705, MSD), targeting eotaxin, eotaxin-3, IL-8, CXCL10, monocyte chemoattractant (MCP)-1/CCL2, MCP-4/CCL13, macrophage derived chemokine (MDC)/CCL22, macrophage inflammatory protein (MIP)-1α, MIP-1β and thymus and activation-regulated chemokine (TARC)/CCL17. Cell free medium was measured as background to subtract from measured sample levels. Panel was run according to manufacturer protocol and electrochemiluminescence was measured on MESO QuickPlex SQ 120MM.

### Immunoblotting

Cells were collected for protein analysis. Protein extraction, concentration measurement and immunoblotting were performed as previously described [25]. SuperSignal Western Blot Enhancer (ThermoFisher) was used for selective antibodies. Immunoblots were visualized by chemiluminescence with SuperSignal Vest Pico Plus (ThermoFisher) on the Cytiva Amersham ImageQuant 800 instrument. List of antibodies and dilutions can be found in Supplementary Table 3. Immunoblots were quantified using ImageJ Fiji version 1.49, representative full-membrane immunoblotting images can be found in Supplementary Figure 6 and 7.

### Gene Expression and Real-time qPCR

Cells for RNA measurement were lysed in Qiazol (Sigma), whereafter RNA was purified using the Direct-zol RNA miniprep kit (Zymo Research) according to manufacturer protocol. cDNA was synthesized from isolated RNA using iScript cDNA Synthesis Kit (Bio-Rad). Real-time qPCR was performed using Taqman Assay and Gene expression Master Mix (Applied Biosystems) and measured on QuantStudio 7 pro (ThermoFisher). Relative gene expression was analysed by 2^-ΔΔCt^ method with normalization to the housekeeping gene *PPIA* [26].

### RNA sequencing

Purified RNA from SC-islets was sent for RNA sequencing at Department of Genomic Medicine at Rigshospitalet Copenhagen University Hospital. RNA was sequenced using the Illumina Stranded Total RNA preparation kit with the Ribo-Zero protocol on an Illumina NovaSeq 6000 platform. Raw count values from in-house primary islets and EndoC-βH5 cells [8] were included and processed alongside SC-islets. Reads were mapped to the GRCh38 v107 Ensembl *Homo sapiens* reference genome using STAR v2.7.11b [27]. EdgeR v4.8.2 with the quasi-likelihood F-test was used for differential expression analysis [28], *p <* 0.05 was considered significant after adjusting with the Benjamini-Hochberg method [29]. pTPM (pseudo-Transcripts Per Million) values were calculated with TMM-normalized counts using edgeR. Concordance correlation coefficients, deming regression and the generalized coefficient of determination (R^2^_G_) were calculated using log10(1+pTPM) [30, 31]. For volcano plots and 2D plots, genes with pTPM < 0.5, in at least four samples, were filtered. Heatmaps were generated using ComplexHeatmap v2.26.0, with per-gene Z-scores calculated from pTPM values of groups within a single heatmap [32]. Pathway analysis was conducted using fGSEA[33], where normalized pathway enrichment scores (NES) were calculated using the MSigDB Reactome database [34–36].

### Statistical analysis

Graphs were generated in Graphpad Prism (version 10.1) and R Studio (version 2025.05.0). Assuming normality, statistical analysis was performed using two-tailed paired or unpaired t-test where appropriate, with no correction for multiple comparison. Raw data values where in some cases Log10.transformed before statistical evaluation. A p-value of <0.05 was considered statistically significant.

## Results

### SC-islets display relevant hormone and transcription factor expression

SC-islets were generated following a 7-stage differentiation protocol (Figure 1A) [18, 24]. Immunofluorescence staining revealed SC-islet cells expressing insulin, glucagon and somatostatin yielding on average 42.4±5.4% INS+ beta cells, 14.3±4.3% GCG+ alpha cells and 1.1±0.7% SST+ delta cells (Figure 1B and 1C). Immature polyhormonal cells accounted for 8.4±6.9% of the cell population, and 33.8±9.9% were undefined (INS-/GCG-/SST-) or off-target cells. Immunoblotting of key islet transcription factors showed a steady expression of Pancreatic Duodenal Homeobox (PDX)-1, NK6 Homeobox (NKX6)-1, and Aristaless-related Homeobox (ARX) across differentiations (Figure 1D).

### SC-islets have a reliable arginine response but inconsistent glucose response

SC-islets were examined for glucose- and arginine-regulated hormone secretion. Insulin secretion in response to high glucose (16.8 mM) was inconsistent, exhibiting batch-dependent responses (Figure 1E). This, however, was comparable to primary islets where a large degree of donor-variation in glucose-induced insulin secretion was observed (Figure 1F). L-arginine consistently stimulated insulin secretion in SC-islets (Figure 1E) but with high variation in primary islets (Figure 1F). EndoC-βH5 cells showed robust insulin secretion at high glucose, but only a modest yet consistent response to L-arginine (Figure 1G).

Glucagon secretion at low or high glucose was inconsistent between batches in both SC-islets and primary islets (Figure 1H+I). Although highly varying, L-arginine stimulated glucagon secretion in 6 out of 7 SC-islet batches and in 3 out of 4 primary islet batches. The hormone content was comparable in SC-islets and primary islets (Figure 1J+K).

### Islet-cell identity gene expression in SC-islets correlates well with primary islets and EndoC-βH5

We next compared the expression of endocrine cell identity genes in SC-islets with that of primary islets and EndoC-βH5 cells. For this purpose, we used in-house RNA-seq data from all models and cell identity genes curated from [37, 38]. Comparison of islet-cell identity genes revealed good concordance of gene expression between SC-islets and primary islets or EndoC-βH5 cells, with concordance correlation coefficients (CCC) of 0.735 and 0.727, respectively (Figure 1L) The CCC between primary islets and EndoC-βH5 cells was 0.665. The expression of alpha-cell identity genes was highly correlated for SC-islets and primary islets with a CCC of 0.861 (Figure 1M), whereas the correlation strength of beta-cell identity gene expression was lower (CCC = 0.585) (Figure 1N). Notably, EndoC-βH5 cells demonstrated comparable correlations with whole-islet identity genes as well as with segregated alpha- or beta-cell identity genes, consistent with patterns observed in both SC-islets and primary islets (Figure 1L-N).

### SC-islets exhibit dose-dependent cytokine responses for cell death and signal transduction

Cytokine (IFNγ+TNFα+IL-1β) exposure for 48h significantly increased SC-islet cell death and caspase 3/7 activity at the two highest doses tested (Figure 2A+B). A time-course experiment using the lowest cytokine dose (50 U/mL IFNγ, 50 U/mL TNFα, and 2.5 U/mL IL-1β) showed no effect on cytotoxicity up to 144 h of exposure (Figure 2C). Based on these findings, we selected two doses of cytokines for the remaining study; a high dose of 1000 U/mL IFNγ, 1000 U/mL TNFα, and 50 U/mL IL-1β for 48 h and a low dose of 50 U/mL IFNγ, 50 U/mL TNFα, and 2.5 U/mL IL-1β for 144 h, respectively referred to as HS (high dose, short exposure) and LL (low dose, long exposure). The rationale being that HS would mimic the high directly toxic level of islet inflammation as seen in T1D, whereas LL would mimic a state of chronic low-grade inflammation characteristic of T2D. To assess activation of key signalling pathways in response to a 30 minute-treatment with cytokines. The protein levels of nuclear factor kappa B (NFκB) inhibitor IκBa, phosphorylated c-Jun N-terminal kinase (pJNK) and signal transducer and activator of transcription (pSTAT)-1 (Figure 2D) were evaluated by western blotting. A cytokine dose identical to HS induced significant downregulation of IκBa and significant increase of pJNK and pSTAT1. A cytokine dose identical to LL only caused significant downregulation of IκBa, with no significant effects on pJNK and pSTAT1 levels (Figure 2D). It is well known that cytokines increase the expression of major histocompatibility complex (MHC) class I in human islets allowing for presentation of peptides to immune cells [39]. In SC-islets, both HS and LL strongly upregulated the protein level of MHC class 1 (Figure 2E).

**Figure 2.**
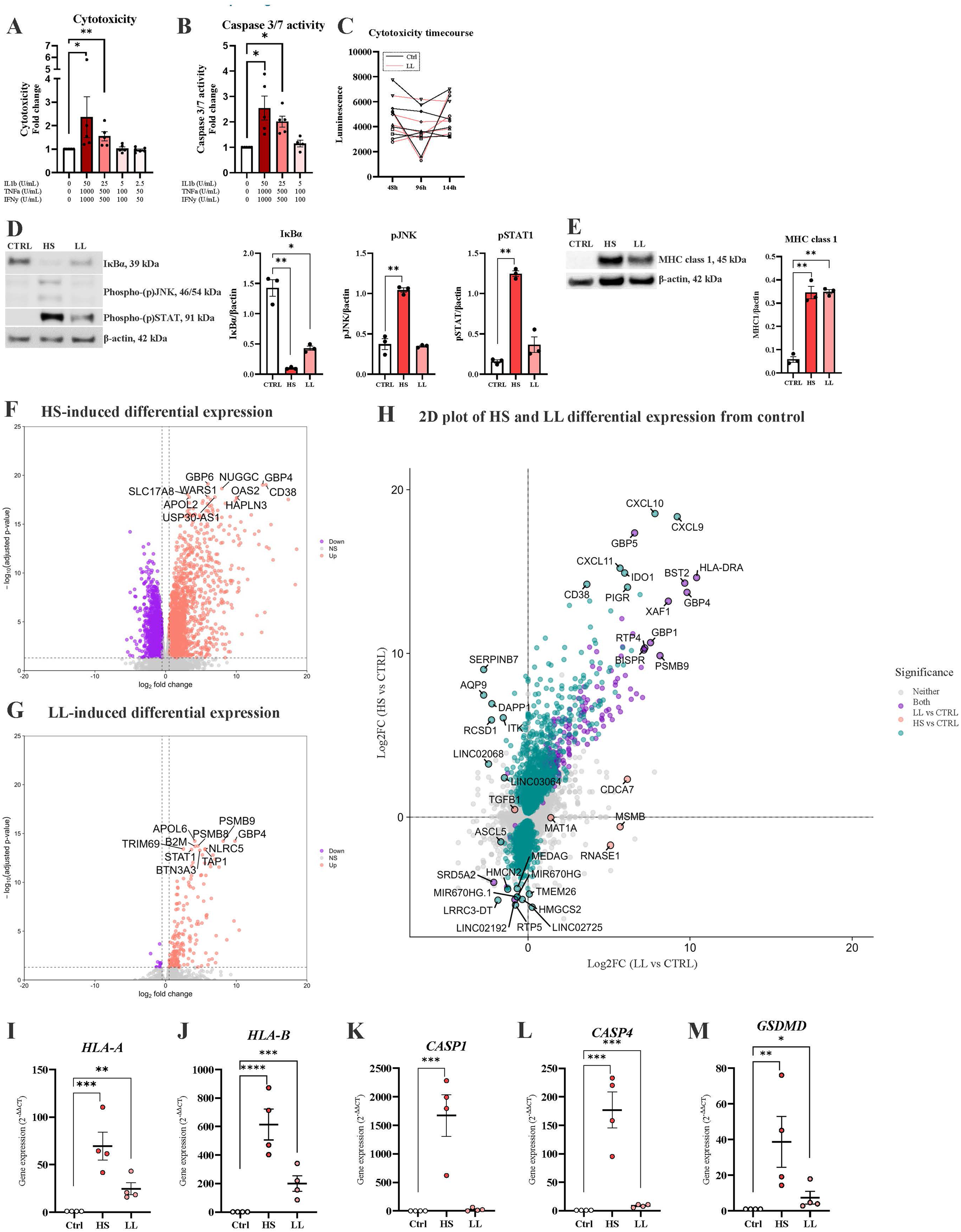
SC-islet cytokine sensitivity to pro-inflammatory cytokine cocktail of IL-1β, TNFα and IFNγ. Cytokine-induced fold change of **A)** Cytotoxicity levels and **B)** Caspase 3/7 activity measured following 48h-stimulation with defined cytokines at different doses. **C)** Time-course for cytotoxicity (luminescence) for low dose stimulation (2.5/50/50) named LL on graph. Remaining graphs are performed with selected doses of HS; 50 U/mL IL-1β, 1000 U/mL TNFα and 1000 U/mL IFNγ and LL; 2.5 U/mL IL-1β, 50 U/mL TNFα and 50 U/mL IFNγ. **D)** Protein collected after 30-minute HS-or LL-stimulation, measured NFκB inhibitor alpha (IκBα), phosphorylated c-Jun N-terminal kinase (pJNK) and phosphorylated signal transducer and translation activator 1 (pSTAT1), normalized to β-actin. **E)** MHC class 1 protein levels following HS for 48h and LL for 144h, normalized to β-actin. Volcano plots for SC-islets differential expression of **F)** HS and **G)** LL, labelled top 10 regulated genes. **H)** 2Dplot of log2(FC) of SC-islets differential expression at HS and LL, HS-only (turquoise), LL-only (orange) or both (purple). Quantitative real-time (q)PCR validation of I) *HLA-A*, J) *HLA-B*, K) *CASP1*, L) *CAPS4* and M) *GSDMD* normalized to housekeeping *PPIA* (2^-ΔΔCT^). All graphs are presented as mean±SEM with individual data points for biological replicates. Paired t-test of log10 (graph A-C), housekeeping-normalized data (D-E) and ΔCT (L-R) graphs A-E and I-M. Data considered significant with p-value <0.05; <0.05*, <0.01**, <0.001***, <0.0001****.

### Cytokine-induced differential gene expression in SC-islets

RNA-seq was performed on SC-islets treated with HS or LL. Differential gene expression (DGE) analysis revealed that while HS (Figure 2F) significantly modulated the expression of 5098 genes in total (2427 upregulated and 2671 downregulated), LL (Figure 2G) had a modest effect on the transcriptome with only 200 genes regulated (190 upregulated and 10 downregulated). Although several genes were similarly regulated across both HS and LL treatment, the impact of the two cytokine conditions on the transcriptome differed markedly (Figure 2H). The top differentially expressed genes under HS and LL are shown in Supplementary Figure 3A. These included genes encoding chemokines and long intergenic non-protein coding genes.

Pathway analysis (Supplementary Figure 3B) revealed shared HS/LL regulation of cytokine signalling and immune recognition, with, not surprisingly, highest induction by HS. Interestingly, five genes were solely regulated by LL; *CDCA7*, *MSMB*, *RNASE1*, *MAT1A,* and *TGFB1*. We selected five key genes for qPCR validation of the RNA-seq data; *HLA-A*, *HLA-B*, *Caspase (CASP)-1, CASP4,* and *Gasdermin D* (*GSDMD*) (Figure 2I-M). The three latter are key genes in pyroptotic cell death, which is an emerging form of programmed cell death that may contribute to beta cell demise in T1D [40]. qPCR of these genes confirmed the RNA-seq findings.

### Cytokine-induced gene expression in SC-islets versus primary islets and EndoC-βH5 cells

We next compared the cytokine-induced transcriptional landscape of SC-islets with that of primary islets and EndoC-βH5 cells. DGE in SC-islets showed considerable overlap with both primary islets and EndoC-βH5 cells (Supplementary Figure 4). Genes associated with programmed cell death, ER stress, autophagy/lysosome, cytokine signalling, islet identity in addition to T1D candidate risk genes, known to play important roles in beta cells and islets, were analysed [6, 8, 41, 42]. Overall, from the heatmaps in Figure 3, the gene expression patterns of genes in these biological terms correlated well between SC-islets primary islets, and EndoC-βH5 cells, albeit with some notable differences. In EndoC-βH5, opposing regulation was seen for *GLIS3, ATF6, TRIB3, GADD45A, RFX6, ISL1, GSDME,* and *BCL2L11* compared to primary islets and SC-islets. In SC-islets, opposing regulation was observed for *IL6R, HHEX, ATG4A, BACH2, PDX1,* and *BID* compared to both primary islets and EndoC-βH5 cells. *TLR4* was upregulated in primary islets and downregulated in both SC-islets and EndoC-βH5. Under LL conditions, SC-islets largely resembled the untreated condition, however, a subset of genes were regulated similarly to the HS condition, including genes involved in immune recognition (e.g. *HLA*) (Supplementary Figure 4A).

**Figure 3.**
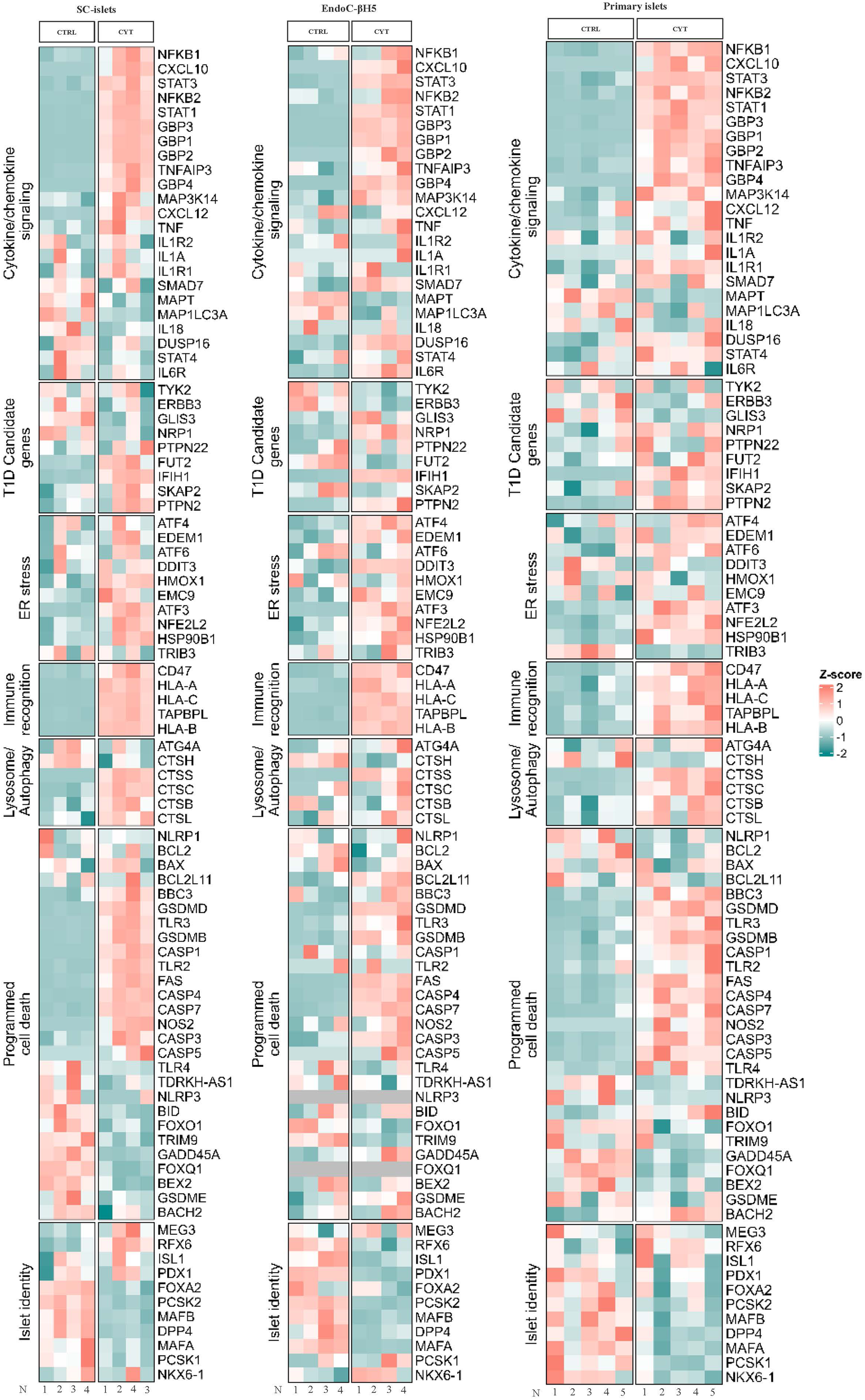
Heatmaps of cytokine-regulated genes commonly associated with programmed cell death (i.e. apoptosis, pyroptosis), endoplasmic reticulum (ER) stress, autophagy/lysosome, cytokine/chemokine signalling, immune recognition, islet identity and diabetes associated risk (T1D and T2D). In-house RNA-seq of A) SC-islets following 48h 50 U/mL IL-1β, 1000 U/mL TNFα and 1000 U/mL IFNγ cytokine-exposure, B) EndoC-βH5 following 24h 50 U/mL IL-1β, 1000 U/mL TNFα and 1000 U/mL IFNγ cytokine-exposure and C) Primary islets following 24h 50 U/mL IL-1β and 1000 U/mL IFNγ cytokine-exposure.

### Cytokine-induced cell death in SC-islets versus primary islets and EndoC-βH5

We performed a head-to-head comparison of HS and LL treatments in SC-islets, primary islets, and EndoC-βH5 cells regarding cytokine toxicity. Cytotoxicity (Figure 4A-B) and caspase-3/7 activity (Figure 4D-E) were increased by HS, but not LL, in SC-islets and primary islets. In contrast, in EndoC-βH5 cells, LL, but not HS, increased cytotoxicity (Figure 4C) and caspase 3/7 activity (Figure 4F). The protein levels for markers of apoptotic and pyroptotic cell death were measured (Figure G-I). Full-length caspase-7 was increased by both HS and LL in SC-islets and primary islets (Figure 4J-L). However, only statistically significantly in primary islets following LL stimulation. Cleavage of caspase-7 was only detected in islets treated with HS (Figure 4M-O). In SC-islets, full-length caspase-3 (Figure 4P) and cleaved caspase-3 (Figure 4R) were upregulated by HS, with no regulation by LL. Full-length GSDMD was upregulated in SC-islets, EndoC-βH5, and primary islets by HS and/or LL (Figure 4T-V), with no change in the level of cleaved GSDMD (Figure 4T). Cleaved GSDMD was undetectable in primary islets and EndoC-βH5 cells (LPS-stimulated THP-1 cells used as positive control) (Figure 4H-I).

**Figure 4.**
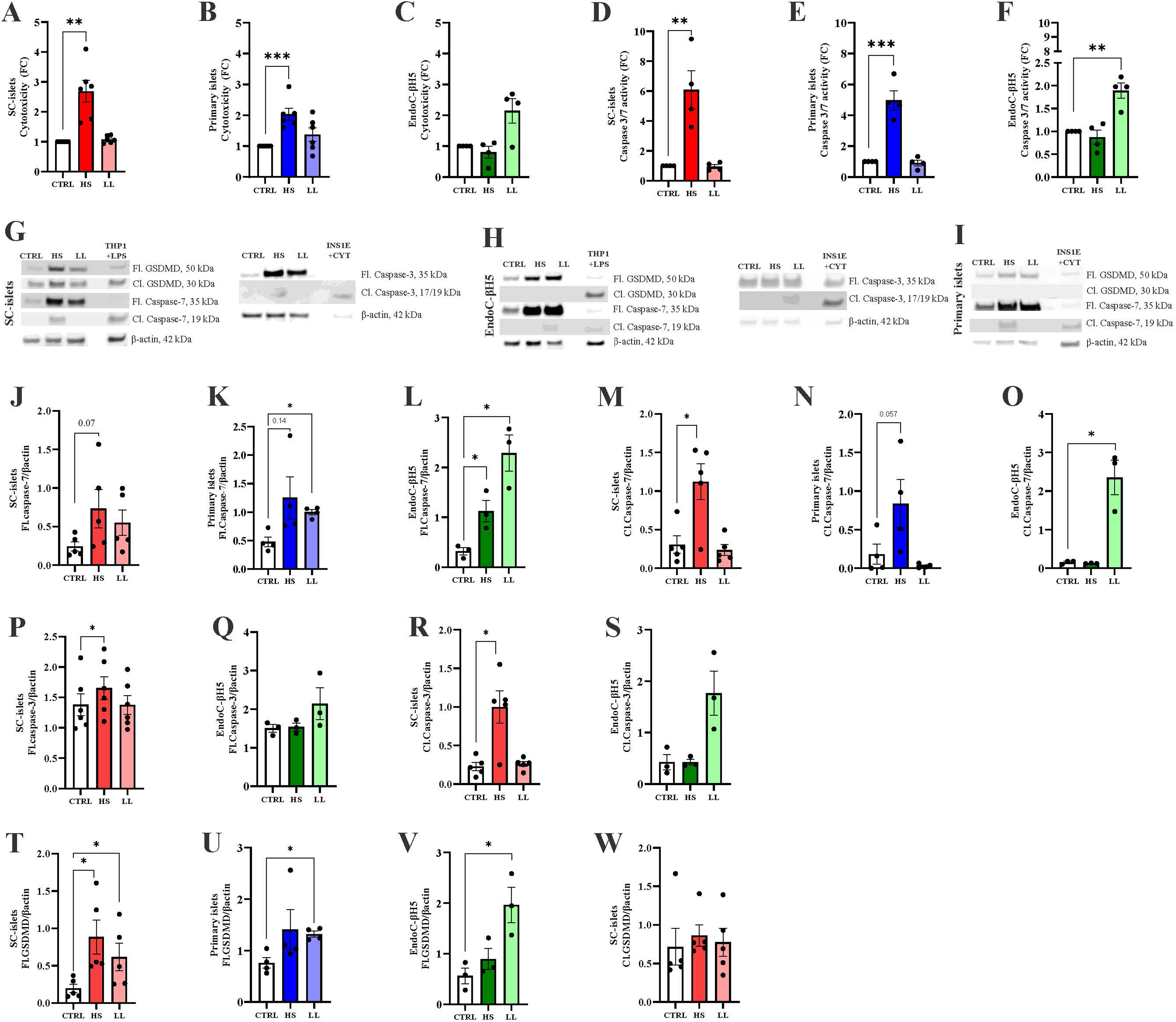
HS- and LL-induced cell death activation of SC-islets (red) with comparison to primary islets (blue) and EndoC-βH5 cells (green). Fold change (FC) **A-C)** cytotoxicity levels and **D-F)** Caspase 3/7 activity. Representative immunoblotting images from G) SC-islets, H) EndoC-βH5 cells, and I) Primary islets. Protein levels of **L-L)** cleaved caspase-7, **M-O)** full-length caspase-7, **P-Q)** cleaved caspase-3, **R-S)** full-length caspase-3, **T-V)** full-length GSDMD and **W)** cleaved GSDMD. Protein expression normalized to β-actin expression for all models. **U)** GSDMD representative western blot images, with LPS-treated THP-1 monocytes or cytokine exposed INS-1E as loading control. All graphs are presented as mean±SEM with individual data points for biological replicate or donor measurements. Statistical analysis by paired t-test on log10-transformed (graph A-F) or raw/normalized data (graph J-W), with comparison to control, 2 comparisons in graph A-F+J-W. Data considered significant with p-value <0.05: *<0.05, **<0.01, ***<0.001.

### Cytokine-mediated chemokine secretion in SC-islets versus primary islets

Islets and beta cells secrete chemokines in response cytokine exposure [8, 9]. We therefore examined chemokine secretion from SC-islets and primary islets following HS and LL exposure using a 10-plex chemokine panel. A high degree of similarity in chemokines secretion were seen between the two islet systems, with HS inducing a significant increase in all chemokines measured (Figure 5A-J). In contrast, LL-induced chemokine secretion was less pronounced, with significant increases observed for eotaxin, CXCL10, CCL22, MIP-1β (Figure 5A+D+I), and CCL17 in both islet models (Figure 5J).

**Figure 5.**
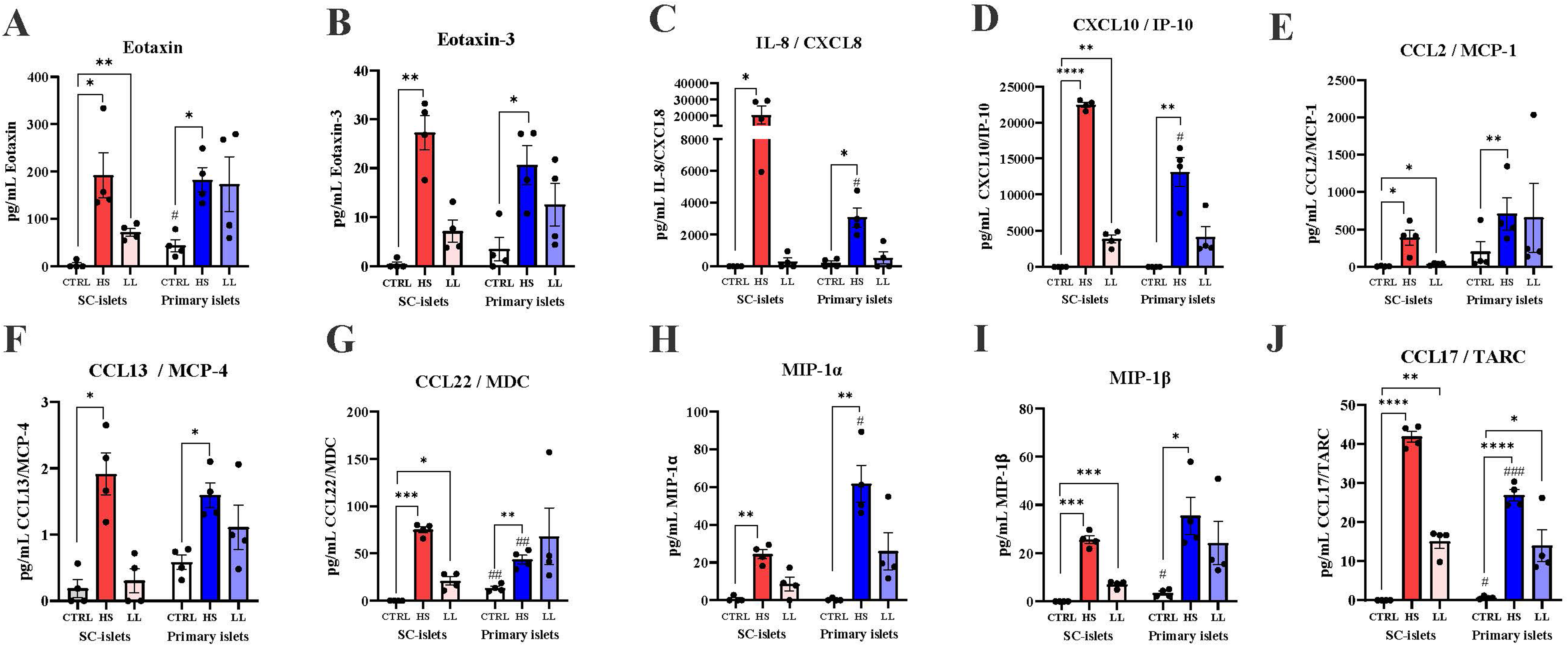
Cytokine-regulated chemokine secretion of SC-islets (red) and primary islets (blue). Chemokine levels (pg/mL), for **A)** Eotaxin, **B)** Eotaxin-3, **C)** IL-8/CXCL8, **D)** IFNγ-induced protein-10 (IP-10/CXCL10), **E)** Monocyte Chemoattractant (MCP)-1/CCL2, **F)** MCP-4/CCL13, **G)** Macrophage derived chemokine (MDC)/CCL22, **H)** Macrophage Inflammatory Protein (MIP)-1α, **I)** MIP-1β and **J)** Thymus and Activation-regulated chemokine (TARC)/CCL17. Values below range were considered 0 pg/mL. All graphs are presented as mean±SEM with individual biological replicate or donor measurements. Statistical analysis by paired t-test on raw data with comparison to sample control (significance marked with *) and unpaired between islet models (significance marked with #), total 7 comparisons. Data considered significant with p-value <0.05: */#<0.05, **/##<0.01, ***/###<0.001, ****<0.0001.

### Cytokine-induced modulation of hormone secretion in SC-islets versus primary islets and EndoC-βH5

SC-islets, primary islets, and EndoC-βH5 cells were assessed for functional glucose- and arginine-stimulated hormone secretion following HS and LL treatment. HS, but not LL, significantly increased insulin secretion at low glucose (2.8 mM) in SC-islets (Figure 6A). The L-arginine-induced insulin response was not affected by HS or LL. In primary islets, insulin secretion at high glucose (16.8 mM) was increased by both HS and LL (Figure 6B), however, only statistically significant following LL stimulation. In EndoC-βH5 cells, the strong glucose-induced insulin secretion was decreased in response to cytokines (Figure 6C), with LL exerting the strongest inhibition. Additionally, the L-arginine-induced insulin secretion increased following HS exposure.

**Figure 6.**
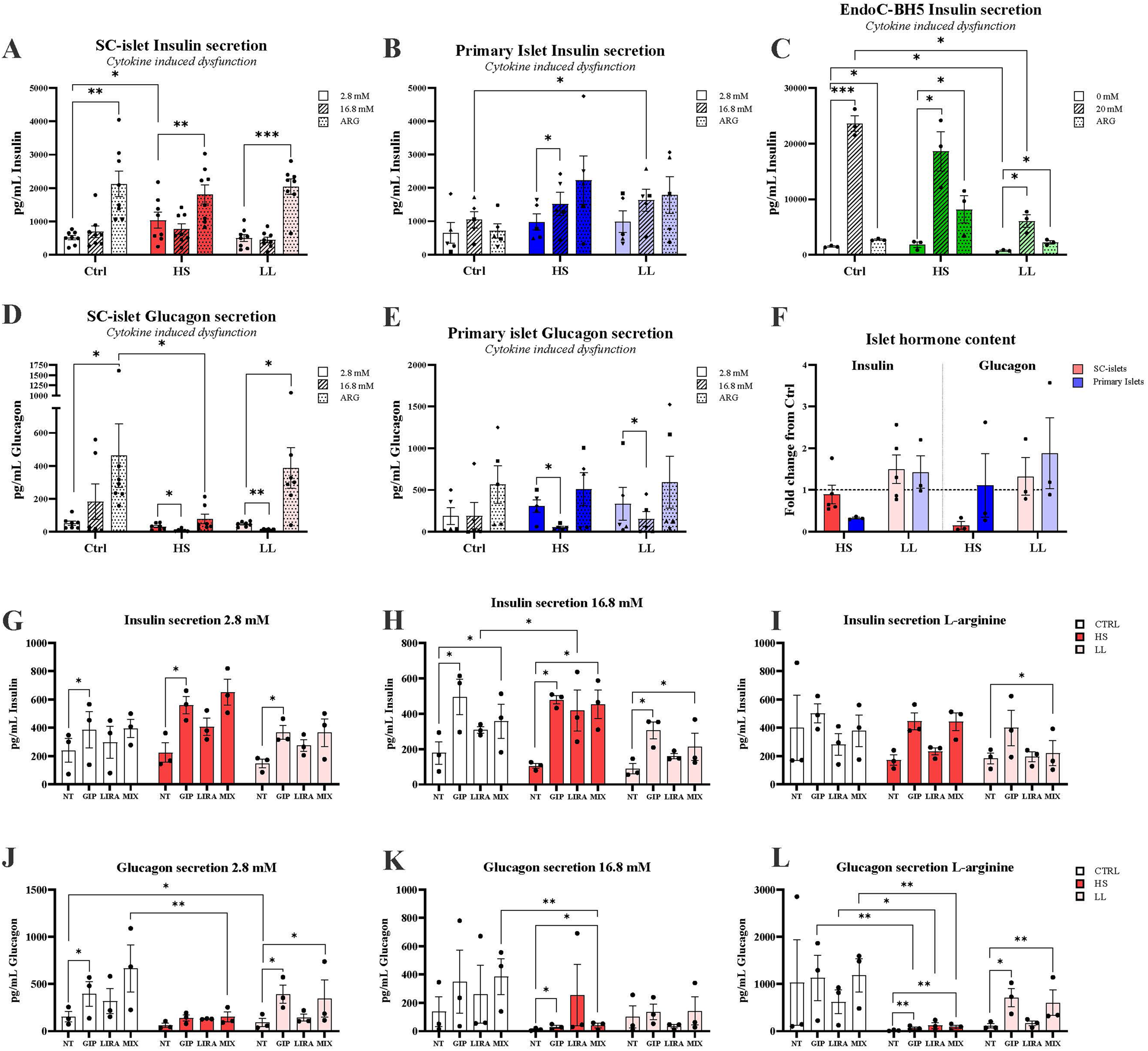
Cytokine-induced dysfunction following HS and LL in SC-islets (red), Primary islets (blue) and EndoC-βH5 cells (green). Insulin secretion following nutrient stimulated secretion assay using 0-2.8 mM glucose as low (clear bar), 16.8-20 mM glucose as high (striped bar) with 15 mM L-Arginine (dotted bar), in **A)** SC-islets, **B)** Primary islets and **C)** EndoC-βH5. Glucagon secretion following HS and LL was measured in **D)** SC-islets and **E)** Primary islets. **F)** Fold change of insulin and glucagon content measured in SC-islets (red) and Primary islets (blue), with stripped line at 1 as control fold. SC-islets hormone secretion assay with acute incretin treatment of 1 uM GIP, 1 uM GLP-1 (Liraglutide) or in combination “MIX”. Incretin regulated insulin secretion at **G)** 2.8 mM glucose, **H)** 16.8 mM glucose and **I)** with 15 mM L-arginine, and glucagon secretion at **J)** 2.8 mM glucose, **K)** 16.8 mM glucose and **L)** with 15 mM L-arginine. Data A-L presented as pg/mL, mean±SEM with individual biological replicates or donor measurements. Paired t-test comparing Log10-transformed data was performed with 12 comparisons on graphs A-E, 4 comparisons performed on graph F and 17 comparisons on graph G-L. Data considered significant with p-value <0.05: *<0.05, **<0.01, ***<0.001.

Regarding glucagon secretion, HS, but not LL, significantly suppressed L-arginine induced glucagon release from SC-islets (Figure 6D). Furthermore, both HS and LL lowered glucagon secretion at high, but not low, glucose in SC-islets. A similar effect, namely reduced glucagon secretion at high glucose, was seen in primary islets (Figure 6E). Analysis of hormone content revealed that HS decreased insulin in primary islets, but not in SC islets (Figure 6F). Contrary to this, HS decreased the glucagon content in SC islets, but not in primary islets, though a high degree of variation was observed (Figure 6F). LL treatment did not affect hormone content in either model.

### Incretin-stimulated hormone secretion in cytokine exposed SC-islets

Finally, based on our recent findings that the incretin hormones can improve the secretory function of cytokine-exposed human islet microtissues [23], we examined the acute effects of glucose-dependent insulinotropic polypeptide (GIP) or Liraglutide (LIRA) (a long-acting GLP-1 receptor agonist) or a combination of both (MIX) on SC islets following HS or LL treatment. GIP elevated insulin secretion at both 2.8 mM (Figure 6G) and 16.8 mM glucose (Figure 6H) in all conditions but had no effect on L-arginine-induced insulin secretion (Figure 6I). MIX was additionally shown to increase insulin secretion at 16.8 mM glucose. GIP alone or in combination (MIX) elevated glucagon secretion at 2.8 mM glucose (Figure 6J) and L-arginine (Figure 6L) for control and LL. The incretin treatments did not restore glucagon secretion in HS (Figure 6J-L), however, increased glucagon secretion at 16.8 mM (Figure 6K) and L-arginine (Figure 6L). Liraglutide (LIRA) alone had no effect on SC-islets in any condition.

## Discussion

In this study, we generated SC-islets containing GCG^+^, SST^+^, and INS^+^ cells. We thoroughly investigated the SC-islets’ sensitivity and responsiveness to inflammation by exposing them to a mixture of classical proinflammatory cytokines (IL-1β+IFNγ+TNFα). We performed direct head-to-head comparisons with primary human islets and EndoC-βH5 cells. We focused on two cytokine conditions, i.e. HS and LL, as models of ‘acute’ high islet inflammation as seen in T1D and chronic low-grade inflammation as seen in T2D. Our studies on DGE, cell-death responses, chemokine and hormone secretion, as well as regulation of key proteins involved in immune recognition and cell-death pathways, revealed overall good consistency between SC-islets and primary islets with some differences observed in EndoC-βH5 cells which is not surprising.

SC-islets were generated with a similar cell composition as previously shown [16, 18], with ∼42% beta cells, ∼14% alpha cells and ∼1% delta cells. Primary human islets have been reported comprised of ∼57% beta cells, ∼35% alpha cells and ∼5-10% delta cells [10, 43]. The SC-islets’ endocrine cell stoichiometry thereby differs from the bona fide islet, which may influence paracrine signalling and cell-cell interactions. The SC-islets were shown to consist of 66% endocrine cells, which are expected to drive the observation in this study, however, off-target cells effects cannot be ruled out. We did not perform immunofluorescent staining for ghrelin-expressing (*GHRL*) epsilon cells and pancreatic polypeptide-expressing (*PPY*) gamma cells, however, expression of these markers was observed at the mRNA level. Common off-target cell types include pancreatic exocrine and mesenchymal cells, as we observed expression of ductal marker *KRT19*, mesenchyme marker *COL1A1*, and no expression of acinar marker *CPA1*. Stem-cell-derived enterochromaffin cells (SC-EC) markers *LMX1A*, *MME*, *ADRA2A*, *TPH1*, *FEV* and *SLC18A1* [44] were also present. Of note, *FEV* expression is also defined as a transcription factor in alpha cells [37]. Single-cell mRNA and protein measurements would be highly valuable to confirm SC-islet findings.

We were unable to illustrate glucose-regulated insulin or glucagon secretion in both SC-islets and primary islets. These findings were unexpected for primary islets, however, clearly underline primary islet heterogeneity [1, 12]. The secretory maturity of SC-islets has been achieved through longer maturation periods or in vivo maturation [14, 16]. Thereby, defining an optimal timepoint for investigating function of both the alpha- and beta-cell populations would be relevant to pursue for future investigations. Interestingly, L-arginine strongly stimulated glucagon-secretion in both islet models, which was unattainable in previous SC-alpha cells [14]. Insulin secretion by L-arginine is expected to act through CAT2A (gene *SLC7A2*) [45–47], however, *SLC7A2* is lowly expressed in EndoC-βH5 which could partially explain their low arginine sensitivity. Correlation analysis revealed similar gene expression in SC-islets, primary islets, and EndoC-βH5, where SC-islet genes expressional patterns may reflect lower maturation than that of primary islets. The lower degree of maturity may partially explain the lower secretory capacity, for example by the low expression of *MAFA* linked to secretory maturity [48].

SC-islets responded strongly to the HS condition with increased cytotoxicity, caspase 3/7 activation, activation of key signal transduction factors (NFκB, pSTAT1, pJNK) and MHC class 1 expression, comparable to previous observations [20]. The effects of the LL condition were much more subtle with no effects on cell death, pJNK, and pSTAT1. Remarkably, however, LL induced a similar level of MHC class I protein as HS and was almost equally effective as HS in causing IκB degradation. These observations are overall in good agreement with less potent effects of LL compared to HS on *HLA-A/B*, *CASP1/4*, and *GSDMD* expression. Our DGE analysis of RNA-seq data showed that HS caused profound changes to the SC-islet transcriptome with more than 5000 differentially expressed genes which was in sharp contrast to LL which only altered the expression of 200 genes. Despite this, there was a substantial degree of overlap in the top 15 pathways affected by HS and LL. These included pathways related to immune recognition, programmed cell death, and cytokine signalling. Interestingly, HS and LL both increased expression of *CASP7* and *GSDMD*, whereas *CASP3* was only increased by HS. SC-islets exposed to HS exhibited a large degree of overlap in differential expression with primary islets and EndoC-βH5 cells. In EndoC-βH5, we observed opposing regulation of genes related to islet-cell identity, ER stress, and programmed cell death compared to SC-islets and primary islets. The difference in cytokine-sensitivity of EndoC-βH5 compared to the islet models may be due to the lower complexity from the lack of non-beta cells. *TLR4* expression was downregulated in both EndoC-βH5 and SC-islets but upregulated in primary islets. *TLR4* encodes toll-like receptor (TLR)-4, a pattern recognition receptor (PRR) linked to pathogen-associated molecular pattern recognition and inflammasome activation [49–51]. This may affect these two models’ ability to respond to e.g. lipopolysaccharide from E. Coli, which is commonly used to activate inflammasome pathways [49, 52, 53]. SC-islets were also shown to have differential regulated expression of *PDX1* compared to primary islets and EndoC-βH5 cells. Cytokines are known to stimulate loss of identity and therefore a decrease in expression was expected [54, 55]. However, we observed increased *PDX1* in SC-islets after cytokine exposure. *PDX1* is expressed by pancreatic progenitors, which is lost in alpha cells with fate-specification. SC-islets could thereby be more sensitive to de- or trans-differentiation [56, 57]. For the comparison of DGE between SC-islets and primary islets it is important to note that RNA from the latter was collected after 24 h of cytokine exposure, not 48 h as the SC-islets, and also with a HS cocktail without TNFα, which could explain some of the discrepancies observed [58, 59]. We nonetheless consider our comparisons valid enough to give an estimate of the similarity of impact of inflammation on the transcriptome. LL induced unique regulation of five genes, which roles in diabetes are unexplored. This includes LL-induced upregulation of *RNASE1* encoding RNase 1 responsible for depredating extracellular RNA which is anti-inflammatory [60–63].

At the protein level, SC-islets and primary islets were shown to have similar responses for caspase 3/7 activity, cytotoxicity, and protein expression of full-length GSDMD, and caspase-7. Caspase-3 and -7 are key caspases of apoptosis with overlapping functions, however unique targets are present for both caspases, which may influence apoptotic signalling [64, 65]. It is well established that cytokines induce caspase 3/7-induced apoptosis in islets and beta cells [5], thereby, further demonstrating SC-islets as a valid model for studying inflammation-mediated apoptosis as previously observed for stem-cell-derived human beta cells [21]. Signs of pyroptotic cell death in SC-islets were also observed as we detected cleaved, activated GSDMD. However, cleaved GSDMD was present in both untreated and cytokine-treated SC-islets suggesting a basal rate of pyroptosis which was not detectable in primary islets or EndoC-βH5 cells. Full-length GSDMD protein, on the other hand, was upregulated by cytokines in all three cell models. GSDMD is an executioner of pyroptotic cell death, which was recently proposed to be involved in beta-cell demise in T1D [40, 66–69]. It is possible that cytokine-mediated upregulation of GSDMD in islets/beta cells prime them for GSDMD cleavage and pyroptosis induction by other stimuli, however, this awaits further investigation.

SC-islets secreted significantly higher levels of IL-8, CXCL10, CCL22, and CCL17 compared to primary islets in response to HS cytokine exposure, whereas the primary islets demonstrated higher secretion of CCL22 and CCL17 under control conditions. IL-8, an inflammatory mediator linked to poor graft survival following transplantation, has been shown to be secreted by islets following isolation [70]. CXCL10 is typically secreted in response to IFNγ and acts as a chemoattractant for lymphocytes involved in the immune infiltration associated with T1D and T2D [71]. CCL22 and CCL17 are chemoattractants of Th2 lymphocytes and regulatory T cells, which lower inflammation and may play protective roles in autoimmune diabetes, immune tolerance, and graft survival [72–74]. These chemokines may therefore exert bidirectional effects on inflammation. Whereas high levels of CCL22 and -17 could serve beneficial for SC-islet transplantation, inhibition of CXCL10 and IL-8 could help ensure graft survival. The very similar chemokine secretion pattern in SC-islets and primary islets, underscore SC-islets as a valid model for studying islet–immune system communication..

Cytokines (HS) increased insulin secretion from SC-islets at 2.8 mM glucose, indicating that some of the beta cells become ‘leaky’ in response to cytokines possibly because of compromised membrane integrity due to cell death induction. However, such insulin ‘leakiness’ at low glucose was not observed for primary islets or EndoC-βH5 cells despite similar levels of cell death as SC-islets arguing that it’s unrelated to cell death leaving the underlying mechanism(s) unknown at this point. Remarkably, the primary islets gained an increased sensitivity to glucose- and L-arginine after cytokine exposure, which was also the case for EndoC-βH5 regarding L-arginine. whereas L-arginine-stimulated insulin secretion was unaffected by cytokine treatment. This suggests that despite induction of cell death by HS, the persisting beta cells remain functional. A higher functional capacity was observed in EndoC-βH5 cells, highlighting this model’s importance in functional beta cell investigations. The SC-islets and primary islets showed significant glucose-inhibited glucagon secretion following HS. In SC-islets, L-arginine-stimulated glucagon secretion was lost following HS while preserved at LL, which is comparable to previous finding [3, 23]. We did observe this secretory pattern in primary islets. SC-islets showed increased glucagon and insulin secretion following acute incretin treatment, with the highest effects by GIP alone or in combination with LIRA, whereas LIRA alone had minimal effects. Acute GIP treatment, with or without LIRA, significantly elevated both insulin and glucagon secretion from NT, suggesting a role in combatting the secretory. GIP-induced glucagon secretion was lost in HS at low glucose (2.8 mM), however, with significant induction present at high glucose and L-arginine. Generally, the LL dose allowed for a better stimulatory window for glucagon secretion, whereas insulin illustrated similar secretory patterns in HS and LL. We previously demonstrated that low-dose cytokine treatment induces alpha-cell dysfunction in primary islet microtissues resembling that observed in T1D [23], which we successfully reproduced in SC-islets. The LL-induced alpha-cell dysfunction was not observed in primary islets, which may be an attribute of higher degree of variation of primary islets [75]. Interestingly, we observed the HS and LL dose to introduce glucose inhibitory effects in alpha cells glucagon secretion across differentiations, and in primary islets, while also suppressed or elevating *GCG* expression following HS or LL exposure, respectively (Supplementary Figure 5). Collectively, the hormone secretion data obtained suggest that SC-islets have a robust and consistent L-arginine response, however, respond inconsistently to glucose. Glucagon secretion under low glucose or L-arginine is hampered by cytokine treatment, but the SC-islets retain their capacity to release glucagon upon GIP stimulation under LL.

In summary, our data confirm that SC-islets is a promising and valid tool for islet inflammation research, having overall good comparability with primary islets regarding cell death pathways, chemokine secretion, and gene and protein expression profiles The EndoC-βH5 cells demonstrated rather unique cytokine response signatures to that of both SC-islets and primary islets, underscoring the importance of islet architecture, endocrine cell composition and complexity. Overall, SC-islets represents a strong platform for screening pathways related to cell death, transcriptional and translational alterations, however, additional studies are warranted to further improve differentiation protocols to generate more functional SC-islets and exploit their use in diabetes and inflammation research.

## Abbreviations

ARX: Aristaless-related Homeobox
CASP: caspase
CCC: correlation correspondence coefficient
CCL: CC motif ligand
CXCL: C-X-C motif chemokine Ligand
DGE: Differential gene expression
E8: Essential 8
ER: Endoplasmic Retikulum
GCG: Glucagon
GIP: Glucose-Dependent Insulinotropic peptide
GSDM: Gasdermin
hiFBS: heat inactivated Foetal Bovine Serum
HLA: Human Leukocyte Antigen
HS: High dose cytokines Short stimulation
IFN: Interferon
IκBa: Inhibitor of kappa B
IL: Interleukin
INS: Insulin
IPSC: Induced Pluripotent Stem cell
LIRA: Liraglutide
LL: Low dose cytokines Long stimulation
MCP: Monocyte Chemoattractant
MDC: Macrophage derived chemokine
MHC: Major Histocompatibility Complex
MIP: Macrophage Inflammatory Protein
NFκB: Nuclear Factor kappa B
NKX: NK6 Homeobox
PDX: Pancreatic Duodenal Homeobox
pJNK: phosphorylated c-Jun N-terminal Kinase
pSTAT: phosphorylated Signal Transducer and Activator of Transcription
pTPM: pseudo-Transcripts Per Million
R^2^_G_: Generalized R squared
ROCKi: Rho-associated Kinase inhibitor
SC-islets: Stem-cell-derived Islets
SST: Somatostatin
T1D: Type 1 diabetes
T2D: Type 2 diabetes
TARC: Thymus and Activation-regulated chemokine
TNF: Tumour Necrosis Factor

## Ethics approval and consent

Anonymized human samples, i.e. human IPSC line and primary human donor islets, were acquired for academic use.

## Acknowledgements

Karyotype validation of IPSCs were performed in collaboration with the Kennedy Center, arranged with Eva Pihl and Associate Professor Lisbeth Birk Møller. Confocal microscopy was performed at the Bartholin Institute under the guidance of Postdoctoral researcher Atul Anand. We thank Miriam Cnop, Décio L. Eizirik and Natalie Pachera for their expert training in IPSC maintenance and differentiation.

## Author contributions

CABS and JS conceptualized the study. CABS, AJ, RG, JG collected data for the study. CABS, RG, ABM analysed the data. CABS, TVS, FP, and JS had supervisory roles. TF contributed with resources. All authors contributed with critical scientific input. CABS wrote the first draft of the manuscript, commented and reviewed by all co-authors. All authors approved the final version.

## Data Availability Statement

RNA-seq data from EndoC-βH5 are available at Gene Expression Omnibus (GEO) repository, accession number GSE218735. RNA-seq data from SC-islets and primary human islets are available upon reasonable request.

