## Supplementary figures and tables for "Stem-Cell-Derived Islets as a Model of Human Islet Inflammation: A Comparative Analysis of Pro-inflammatory Cytokine Responses"

**Supplementary figure legends**

**Supplementary Figure 1** Induced pluripotent stem cell (IPSC)-line 1023 was validated prior to initiation of study. **A)** Pluripotency staining of fixed IPSC targeting transcription factors OCT4 and Nanog (green) and surface markers SSEA-4 and TRA-1-60 (red), with nuclei staining by DAPI (blue). Scalebar: 200 nm. **B)** Spontaneous differentiation was allowing in un-specialized medium, following staining of markers of Ectoderm (beta-tubulin, green), Mesoderm (vimentin, green) and Endoderm (SOX17, red), with nuclei staining by DAPI (blue). Scalebar: 200 nm. **C)** IPSC were validated for normal Karyotype, based on number of chromosomes and chromosome pattern, performed by Kennedy Center Glostrup.

**Supplementary Figure 2** Checkpoint validations for differentiation of SC-islets. At end stage 1 – "definitive endoderm", the cells were fixed and stained for SOX17 (red) and OCT4 (green) to confirm endodermal marker and loss of pluripotency. At end stage 4 – "pancreatic progenitors", the cells were fixed and stained for PDX1 (red) and NKX6-1 (green) to confirm formation of pancreatic endoderm lineage. Nuclei staining by DAPI (blue). Scalebar: 200 nm.

**Supplementary Figure 3** Differential expression following cytokine exposure. **A)** SC-islet Log10(pTPM+1) for control (blue), LL (light yellow) and HS (yellow) of top common and individually differentially expressed genes, the five unique LL-regulated genes are highlighted with red squares. **B)** Top pathways regulated in HS and LL, fGSEA Pathway analysis from Reactome database (MsigDB version), plotted normalized enrichment score (NES). **C)** Log10(pTPM+1) of LL-unique gene expression in Primary islet and EndoC-βH5. **D)** PCA plot for SC-islet RNAseq sample variation within groups/conditions, shape codes indicate the unique identifier for individual differentiations. Bar graphs presented as mean±SEM with individual data points of biological replicates. Paired t-test performed for bar graphs. Data considered significant with p-value < 0.05: *<0.05, **<0.01, ***<0.001.

**Supplementary Figure 4** High dose cytokine-induced differential expression in human models. Heatmap for LL-regulated expression in SC-islets (as seen in Figure 3). **B)** Volcano plot of differentially expressed genes (log2FC) in primary islets following IFNγ and IL-1β (HS dose, minus TNFα). **C)** Volcano plot of differentially expressed genes (log2FC) in EndoC-βH5 following HS stimulation IFNγ, IL-1β and TNFα. **D)** Venn Diagram of differential expression overlap between EndoC-βH5 (Turquoise), SC-islets (yellow) and Primary islets (pink). **E)** 2Dplot for Log2FC differential expression to control, comparison of HS-stimulated SC-islets and HS-like-stimulated Primary islets. **F)** 2Dplot for Log2FC differential expression to control, comparison of HS-stimulated SC-islets and EndoC-βH5.

**Supplementary Figure 5** Islet-identity gene and protein expression in SC-islets following cytokine exposure (LL and HS). Relative expression by qPCR normalized to housekeeping *PPIA*, presented as $2^{-\Delta\Delta CT}$, for **A)** *INS*, **B)** *PDX1* and **C)** *GCG*. Protein levels of **D)** ARX and **E)** NeuroD1, normalized to beta-actin. **F)** Heatmap of z-score pTPM comparison of identity genes for islet cells. **G)** Immunofluorescent images of SC-islets, for insulin (green), glucagon (red) and nuclei (blue), scalebar: 50 nm. Graphs presented as mean±SEM with individual biological replicates. Paired t-test comparisons performed to control. Data considered significant with p-value < 0.05: *<0.05, **<0.01.

**Supplementary Figure 6** SC-islet representative full-length immunoblots. Individual membranes marked with protein target next to bands. The notations are the biological replicates run on the membrane, numbers do not correspond between gels. Loading controls: LPS-treated THP-1 monocytes or cytokine-exposed INS-1E. Ladder: MagicMark XP (ThermoFisher) with bandwidth 20-250 kDa

**Supplementary Figure 7** EndoC-βH5 cells and primary islet representative full-length immunoblots. Individual membranes marked with protein target next to bands. The notations are the biological or donor replicates run on the membrane. Loading controls: LPS-treated THP-1 monocytes or cytokine-exposed INS-1E. Ladder: MagicMark XP (ThermoFisher) with bandwidth 20-250 kDa

**Supplementary table 1:** Human primary islet donor specifications

| Islet preparation identification | HP-25063-01 | HP-25073-01 | HP-25113-01 | HP-25135-01 | HP-25150-01 | HP-25163-01 |
|---|---|---|---|---|---|---|
| **Donor age (years)** | 59 | 56 | 56 | 40 | 69 | 62 |
| **Donor sex (F/M)** | M | M | M | F | M | F |
| **Donor BMI (kg/m2)** | 24 | 29.1 | 28.3 | 25.0 | 29.4 | 24.0 |
| **Donor HBA1c (%)** | 5.7 | 5.4 | 5.3 | 5.0 | 5.4 | 5.5 |
| **History of diabetes** | No | No | No | No | No | No |
| **Islet Purity (%)** | 85-90 | 90 | 85-90 | 90 | 95 | 95 |
| **Islets used for** | GSIS, Cytotoxiciy, WB | GSIS, Cytotoxiciy, WB | GSIS, Cytotoxiciy, WB, Caspase-3/7 | GSIS, Cytotoxiciy, WB, Caspase-3/7 | GSIS, Cytotoxiciy, WB, Caspase-3/7 | Cytotoxiciy, WB, Caspase-3/7 |

**Supplementary table 2:** Media components and factors used for SC-islet differentiation, based on protocols of Barsby et. al 2022 and Fantuzzi et. al 2022.

| | Target | Duration | Media change | Factors added | Basal Medium |
|---|---|---|---|---|---|
| **IPSC** | Single cell (day -1) | 1 day | 100% change | ROCK Inhibitor (Stem Cell Tech/Millipore) | Essential 8 Medium |
| **Stage 1** | Definitive Endoderm | Day 0 | 100% daily | 100 ng/mL Activin A (Peprotech)<br>5 uM CHIR99021 (Tocris) | B1: MCDB131 (Gibco)<br>0.5% faf BSA (Sigma)<br>2 mM Glutamax (Gibco)<br>1.5 g/L NaHCO3 (Sigma)<br>10 mM Glucose (Gibco) |
| | | Day 1 | 100% daily | 100 ng/mL Activin A<br>0.5 uM CHIR99021 | B1 |
| | | Day 2 | 100% daily | 100 ng/mL Activin A | B1 |

| | | | | | |
|---|---|---|---|---|---|
| **Stage 2** | Gut tube formation | Day 3-5 | 100% daily | 50 ng/mL FGF7 (Peprotech)<br>0.25 mM Ascorbic Acid (Sigma) | B1 |
| **Stage 3** | Pancreatic Progenitor 1 | Day 6-7 | 100% daily | 50 ng/mL FGF7<br>0.25 mM Ascorbic Acid<br>250 nM SANT1 (Sigma)<br>1 uM Retinoic acid (Sigma)<br>100 nM LDN193189 (Selleckchem)<br>200 nM TPB (Santa Cruz) | B2: MCDB131<br>2% faf BSA<br>2 mM Glutamax<br>2.5 g/L NaHCO3<br>10 mM Glucose |
| **Stage 4** | Pancreatic Progenitor 2 | Day 8-11 | 100% daily | 50 ng/mL FGF7<br>0.25 mM Ascorbic Acid<br>100 ng/mL EGF (Peprotech)<br>10 mM Nicotinamide (Sigma)<br>10 ng/mL Activin A<br>250 nM SANT1<br>100 nM Retinoic acid<br>200 nM LDN193189 | B2 |
| **Stage 5** | Endocrine progenitor | Day 12-15 | 100% change | S5: 250 nM SANT1<br>100 nM Retinoic acid<br>100 nM LDN193189<br>1 uM GC1 (Tocris)<br>100 nM GSiXX (Sigma)<br>10 uM ALK5inh (Repsox, Selleckchem)<br>20 ng/mL Beta Cellulin (Peprotech) | B3: MCDB131<br>2% faf BSA<br>2 mM Glutamax<br>1.5 g/L NaHCO3<br>20 mM Glucose<br>1:200 ITS-X (Gibco)<br>10 ug/mL Heparin<br>10 uM ZnSO4 (Sigma)<br>100 U/mL Penicillin and 100 ug/mL Streptomycin (Gibco) |
| **Stage 6** | Immature Islet Cells | Day 16-23 | 50% every second day | 100 nM LDN193189<br>1 uM GC1<br>100 nM GSiXX<br>10 uM ALK5inh | B3 |
| **Stage7** | Maturing Islet Cells (SC-islets) | Day 24-45 | 50% every second day | 1 mM N-Acetylcysteine (Sigma)<br>10 nM T3 (Sigma)<br>500 nM ZM447439 (Selleckchem) | CMRL1066 (Corning)<br>2g/100mL faf BSA<br>2 mM Glutamax<br>100 U/mL Penicillin and 100 ug/mL Streptomycin<br>1:200 ITS-X<br>0.5 mM pyruvate (Gibco)<br>10 ug/mL heparin<br>10 uM ZnSO4<br>1X Trace element A (Gibco)<br>1X trace element B (Gibco)<br>1X CD lipid (Gibco) |

**Supplementary table 3:** List of primary and secondary antibodies

| Antibody Target (human) | Source species | Cat.nb#, Company | Dilution | |
|---|---|---|---|---|
| | | | Immunofluoresence | Immunoblotting |
| OCT4 | Rabbit | 2840, Cell Signalling | 1:400 | - |
| Nanog | Rabbit | 4903, Cell Signalling | 1:400 | - |
| TRA-1-60 | Mouse | MA1-023, Life technologies | 1:200 | - |
| SSEA4 | Mouse | MA1-021, Life technologies | 1:500 | - |
| SOX17 | Goat | AF1924, R&D bioscience | 1:400 | - |
| PDX1 | Goat | AF2419, R&D bioscience | 1:500 | - |
| | Rabbit | 5679, Cell Signalling | - | 1:500 |
| NKX6-1 | Mouse | 5633022, BD bioscience | 1:250 | 1:500 |
| INS (C-peptide) | Rat | GN-ID4, DSHB | 1:300 | - |
| GCG | Mouse | G2654, Sigma | 1:1000 | - |
| SST | Rabbit | 108456, Abcam | 1:1000 | - |
| Full-length Caspase 3 | Rabbit | PA5-77887, Life technologies | - | 1:1000 |
| Cleaved Caspase 3 | Rabbit | 9661, Cell Signalling | - | 1:500 |
| Full-length Caspase 7 | Mouse | 9494, Cell Signalling | - | 1:1000 |
| Cleaved caspase 7 | Rabbit | 9491, Cell Signalling | - | 1:500 |
| Phospho-STAT1 | Rabbit | 7649, Cell Signalling | - | 1:1000 |
| Phospho-JNK | Rabbit | 9252, Cell Signalling | - | 1:1000 |
| IkBa | Mouse | J1512, Santa Cruz | - | 1:1000 |
| MHC class 1 | Mouse | ALX-805-711-C100, Enzo | - | 1:1000 |
| GSDMD | Rabbit | 69469, Cell Signalling | - | 1:500 |
| Cleaved N-GSDMD | Rabbit | 36425, Cell Signalling | - | 1:500 |
| ARX | Rabbit | PA5-40407, Life technologies | - | 1:500 |
| NeuroD1 | Rabbit | 4373, Cell Signalling | - | 1:1000 |
| Rat-Alexa 488 | Donkey | 712-545-150, Jackson ImmunoResearch | 1:500 | - |
| Goat-TRITC | Donkey | 705-025-147, Jackson ImmunoResearch | 1:500 | - |
| Mouse-FITC | Donkey | 715-095-151, Jackson ImmunoResearch | 1:500 | - |
| Mouse-TRITC | Donkey | 715-025-151, Jackson ImmunoResearch | 1:500 | - |
| Rabbit-FITC | Donkey | 711-095-152, Jackson ImmunoResearch | 1:500 | - |
| Rabbit-TRITC | Donkey | 711-025-152, Jackson ImmunoResearch | 1:500 | - |
| Mouse-Alexa 647 | Donkey | 715-605-150, Jackson ImmunoResearch | 1:500 | |
| HRP-conjugated anti-mouse | - | 7076, Cell Signalling | - | 1:1000 |
| HRP-conjugated anti-rabbit | - | 7074, Cell Signalling | - | 1:2000 |

### *SUPPLEMENTARY FIGURE 1*

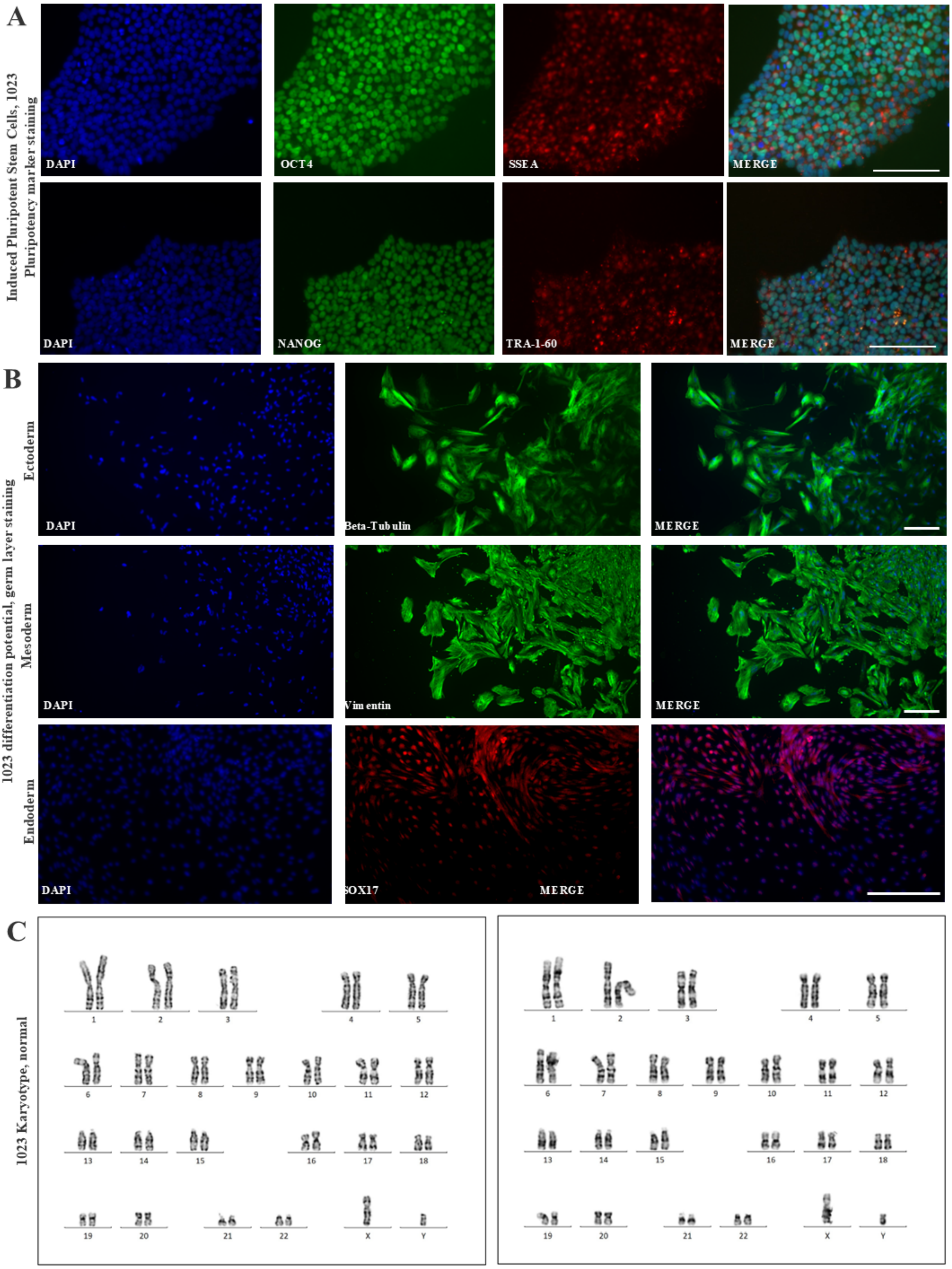

### SUPPLEMENTARY FIGURE 2

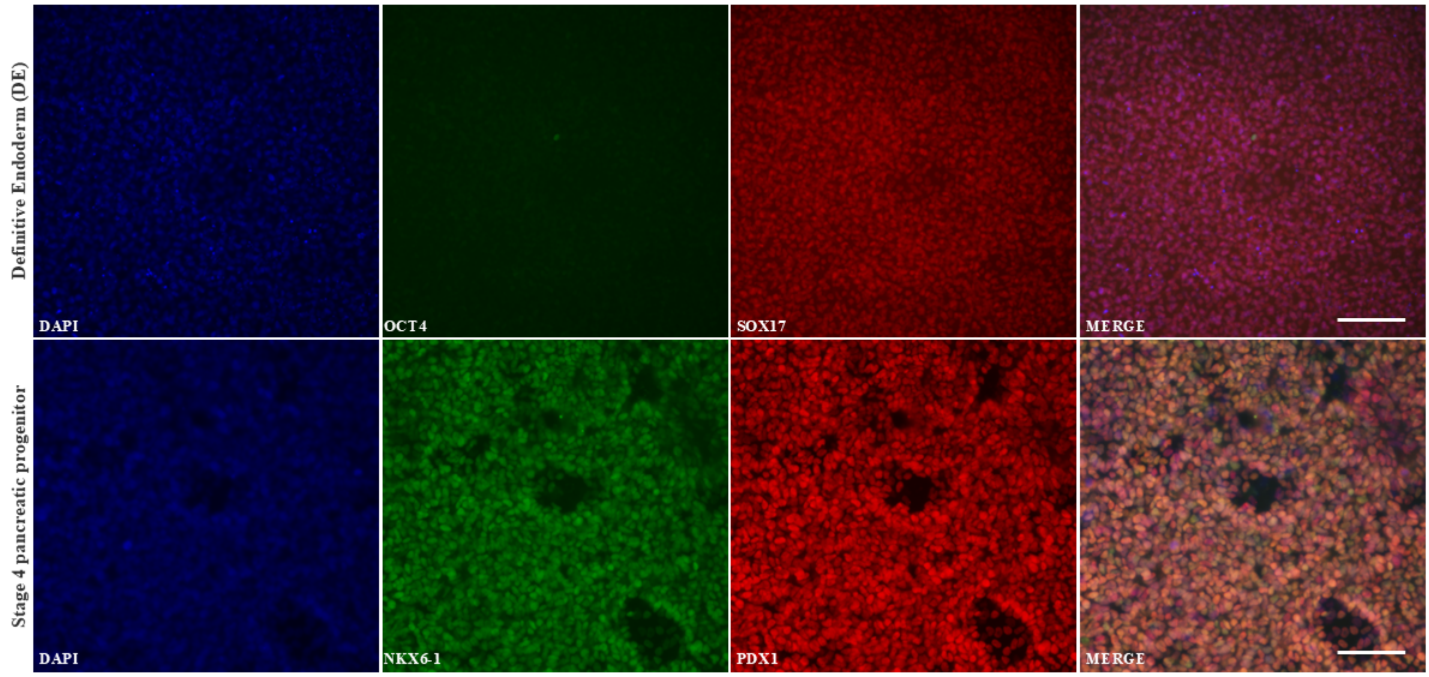

### SUPPLEMENTARY FIGURE 3

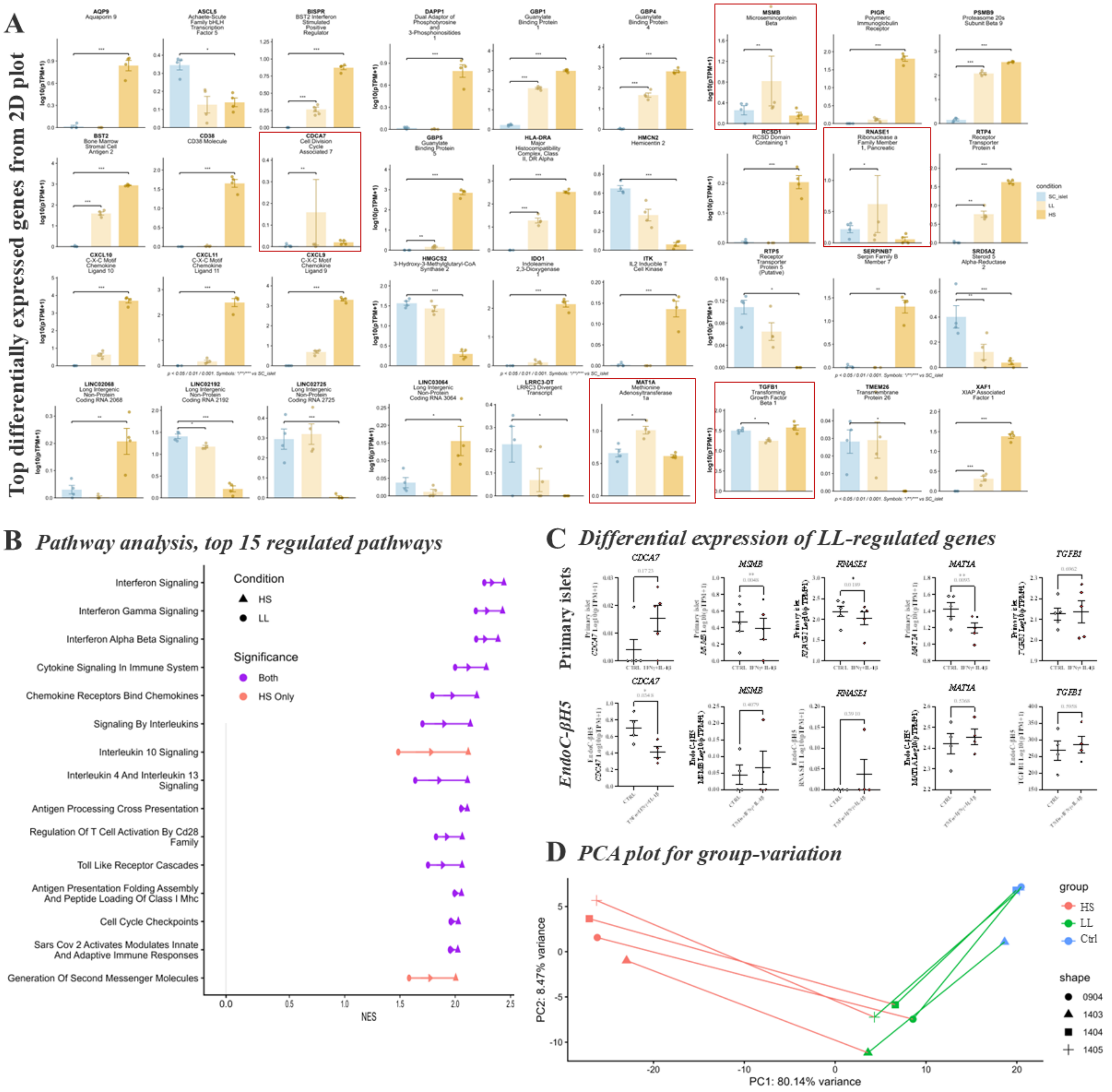

### *SUPPLEMENTARY FIGURE 4*

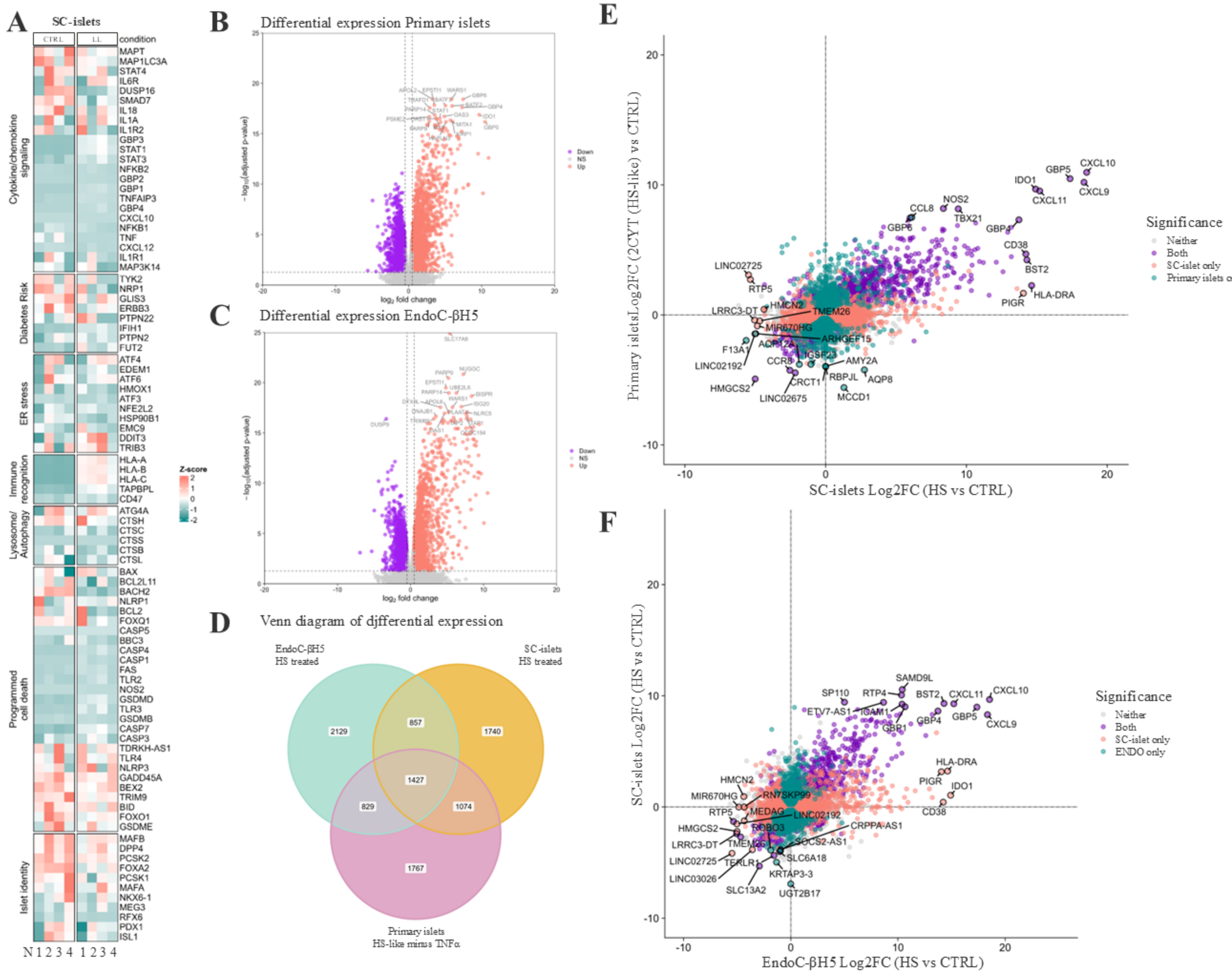

#### *SUPPLEMENTARY FIGURE 5*

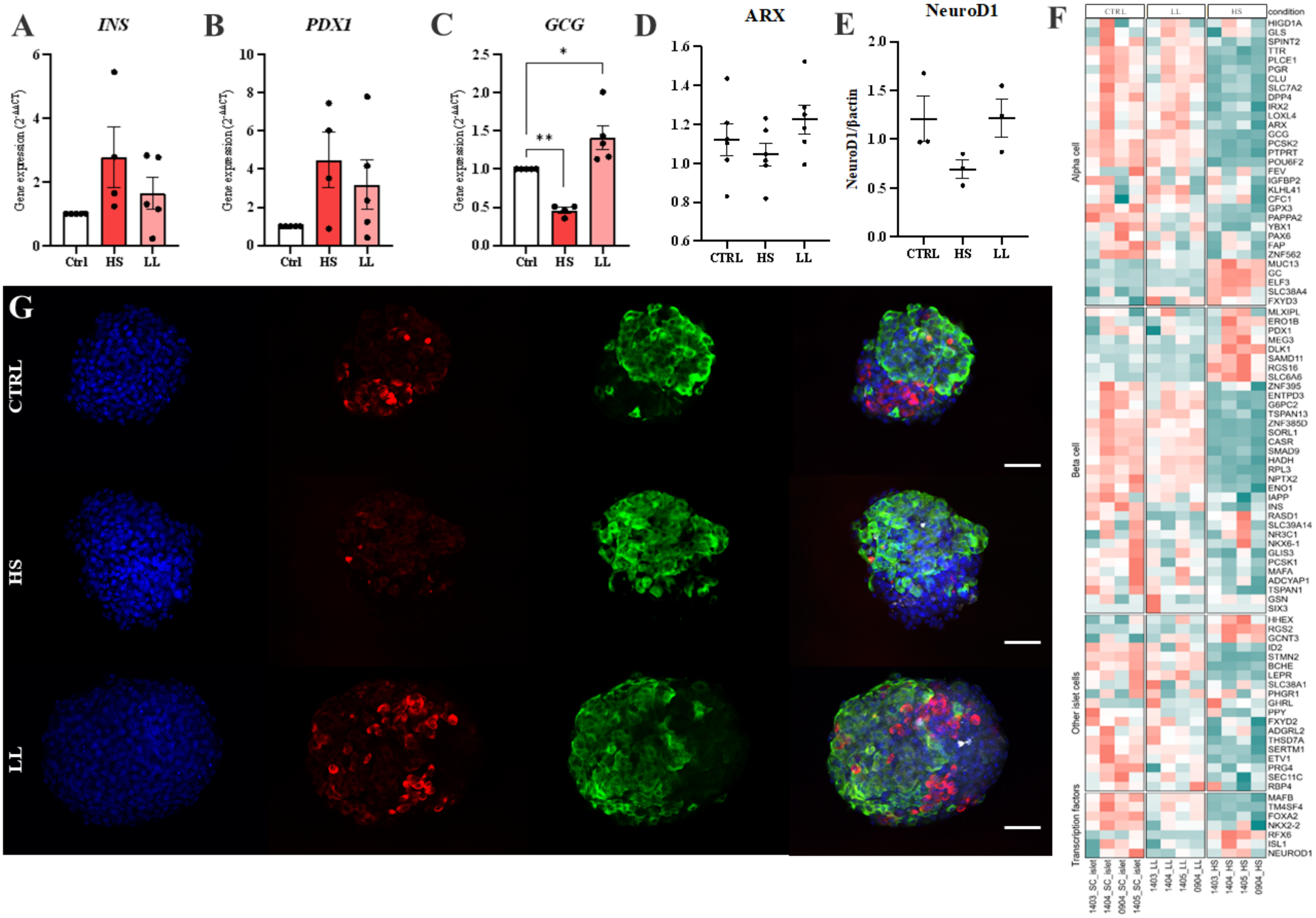

### SUPPLEMENTARY FIGURE 6

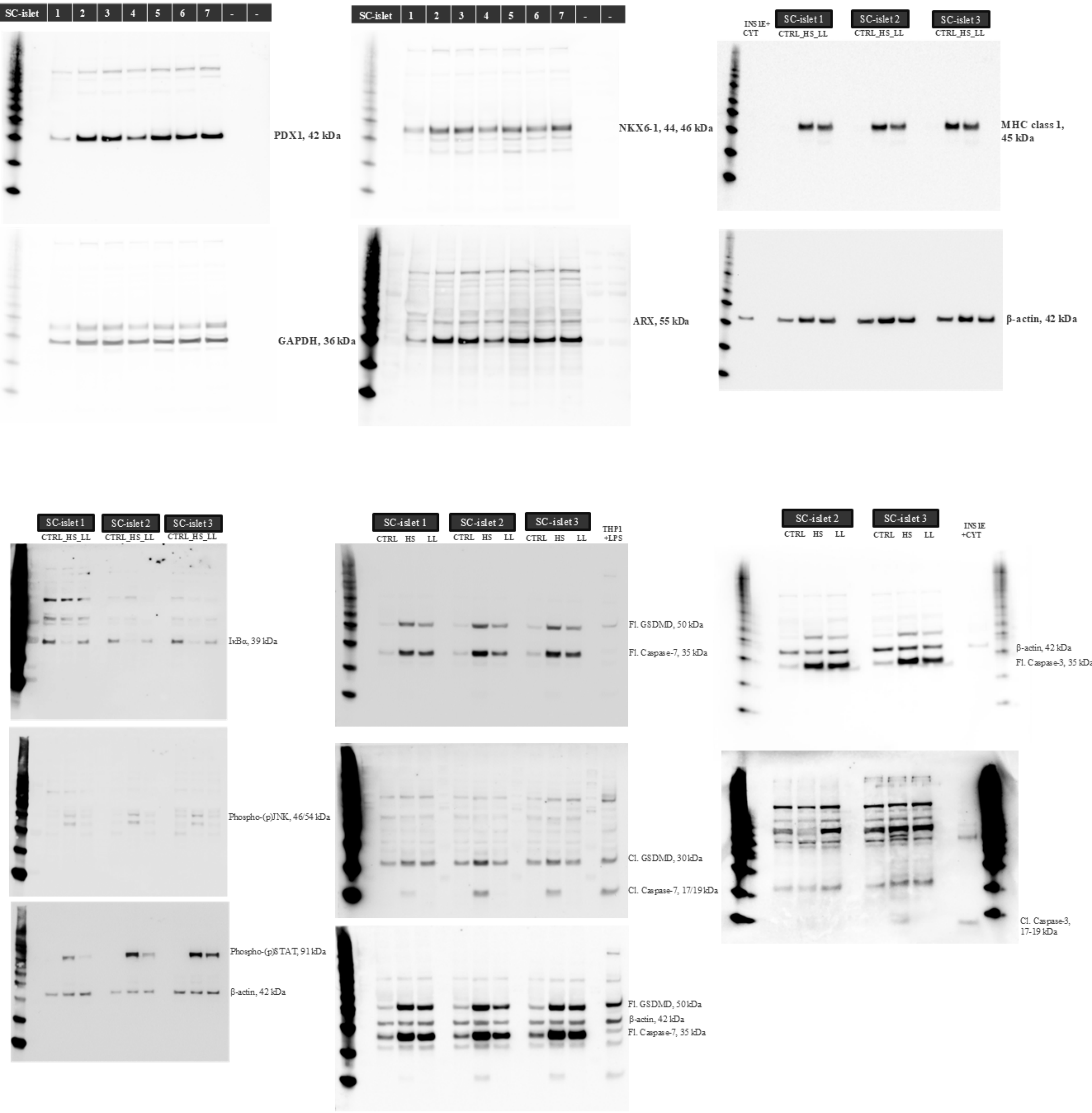

### *SUPPLEMENTARY FIGURE 7*

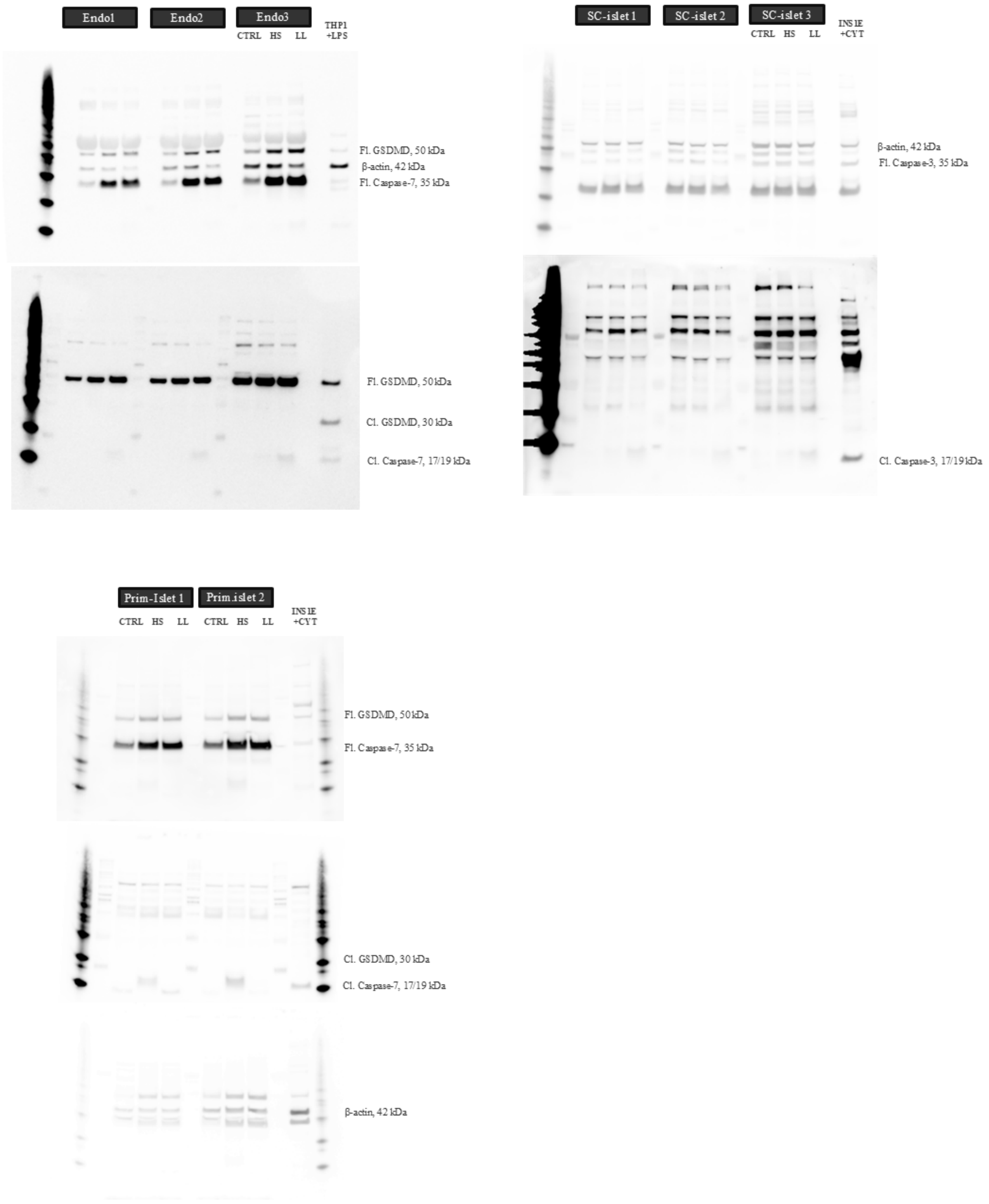
